# Basic helix-loop-helix pioneer factors interact with the histone octamer to invade nucleosomes and generate nucleosome depleted regions

**DOI:** 10.1101/2022.03.17.484790

**Authors:** Benjamin T. Donovan, Hengye Chen, Priit Eek, Zhiyuan Meng, Caroline Jipa, Song Tan, Lu Bai, Michael G. Poirier

**Affiliations:** Biophysics Graduate Program, The Ohio State University, Columbus, OH 43210, USA; Department of Biochemistry and Molecular Biology, Center for Eukaryotic Gene Regulation, The Pennsylvania State University, University Park, PA 16802, USA; Department of Physics, The Ohio State University, Columbus, OH 43210, USA; Huck Institutes of the Life Sciences, The Pennsylvania State University, University Park, PA 16802, USA; Department of Physics, The Pennsylvania State University, University Park, PA 16802, USA; Department of Chemistry & Biochemistry, The Ohio State University, Columbus, OH 43210, USA

**Keywords:** gene regulation, chromatin biology, pioneer transcription factors, dissociation rate compensation mechanism, nucleosome depleted regions, single molecule measurement, cryoelectron microscopy single particle analysis

## Abstract

Nucleosomes drastically limit transcription factor (TF) occupancy, while pioneer transcription factors (PFs) somehow circumvent this nucleosome barrier. In this study, we compare nucleosome binding of two conserved *S. cerevisiae* basic helix-loop-helix (bHLH) TFs, Cbf1 and Pho4. A Cryo-EM structure of Cbf1 in complex with the nucleosome reveals that the Cbf1 HLH region can electrostatically interact with exposed histone residues within a partially unwrapped nucleosome. Single molecule fluorescence studies show that the Cbf1 but not the Pho4 HLH region facilitates efficient nucleosome invasion by slowing its dissociation rate relative to DNA through interactions with histones. *In vivo* studies show that this enhanced binding provided by the Cbf1 HLH region enables nucleosome invasion and ensuing repositioning. These structural, single molecule, and *in vivo* studies reveal the mechanistic basis of dissociation rate compensation by pioneer factors and how this translates to facilitating chromatin opening inside cells.

## INTRODUCTION

Chromatin structure regulates occupancy of DNA binding proteins genome wide. A primary example is that nucleosome positioning relative to a transcription factor (TF) binding site can regulate TF occupancy by orders of magnitude (Adams and Workman, 1995; Li and Widom, 2004; Luo et al., 2014a; Polach and Widom, 1995), which is crucial to gene regulation (Bai and Morozov, 2010; Bai et al., 2010; Dadiani et al., 2013; Donovan et al., 2019a; Rodriguez and Larson, 2020). However, DNA binding proteins use a myriad of strategies to overcome barriers presented by the nucleosome. Binding upon transient partial nucleosome unwrapping is a primary mechanism by which TFs gain access to sites within nucleosomes (Li and Widom, 2004; Li et al., 2005; Polach and Widom, 1995; Tims et al., 2011). This site exposure mechanism predicts a reduction in TF *binding* based on the probability the DNA target site is exposed (North et al., 2012). More recently, single molecule studies have revealed that nucleosomes also regulate TF occupancy through influencing TF *dissociation*. Nucleosomes can reduce TF occupancy through accelerated dissociation (Chen and Bundschuh, 2014; Luo et al., 2014a) and can enhance occupancy by slowed dissociation (Cirillo and Zaret, 1999; Donovan et al., 2019b).

Pioneer factors (PFs) are key transcription factors that bind their target sites within nucleosomes as efficiently as naked DNA (Soufi et al., 2014; Zaret and Carroll, 2011). They initiate the formation of nucleosome depleted regions (NDRs) (Bai et al., 2011; Heinz et al., 2010), and often function as master regulators of cell fate (Holtzinger and Evans, 2006; Lee et al., 2005). For some PFs, slowed dissociation from nucleosomes is an important property (Cirillo and Zaret, 1999; Donovan et al., 2019b; Makowski et al., 2020) because it enhances the effective affinity of PFs to target sites within nucleosomes despite the limited binding site accessibility. Such “dissociation rate compensation mechanism” may enable, or at least facilitate, these factors to target nucleosomal sites so that they can initiate the formation of NDRs and chromatin opening (Yan et al., 2018; Zaret and Carroll, 2011). However, the mechanisms underlying the nucleosome-dependent slow dissociation and how essential this PF binding property is for the establishment of NDRs *in vivo* is unknown.

Cbf1, a basic Helix Loop Helix Leucine Zipper (bHLHZ) transcription factor, can function as a pioneer factor and generate NDRs near its binding sites in budding yeast (Yan et al., 2018). Using *in vitro* single molecule assays, we recently showed that Cbf1 can access transiently exposed binding sites within partially unwrapped nucleosomes to stabilize the unwrapped state without histone eviction. These results are consistent with recent structural studies that report PFs bound to nucleosomes stabilize distorted (Dodonova et al., 2020) and even partially unwrapped DNA (Michael et al., 2020). Cbf1 binding to partially unwrapped nucleosomes results in an ∼100-fold reduction in binding rate, however the roughly equal (∼30 fold) reduction in dissociation rate enables Cbf1 to bind nucleosomes with similar affinities as to naked DNA (Donovan et al., 2019b). This finding strongly suggests that Cbf1 has additional interactions with nucleosomes compared to naked DNA. However, the molecular basis of these nucleosome specific interactions is unknown.

Cbf1 is commonly contrasted with Pho4, another bHLH TF from budding yeast because of their highly similar DNA binding domains and subsequent preference for E-box motifs. DNA binding mechanisms for both proteins have been extensively characterized (Cave et al., 2000; Maerkl and Quake, 2007; Shimizu et al., 1997; Wieland et al., 2001). Interestingly, unlike Cbf1, nucleosomes almost completely block Pho4 binding (Zhou and O’Shea, 2011). It remains unknown how two similar TFs recognize nearly the same DNA target sequence despite such drastic differences in binding behaviors to nucleosomes.

In this manuscript, we find major differences in how nucleosomes regulate occupancy of Pho4 and Cbf1; Cbf1 undergoes slowed dissociation from nucleosomes while Pho4 does not. Interestingly, while both proteins are significantly disordered, only the structured regions are required for DNA and nucleosome binding. This result is confirmed by our Cryo-EM structure of Cbf1 in complex with the nucleosome, which demonstrates that Cbf1 traps the nucleosome in a partially unwrapped state and reveals that the structured C-terminal HLH region can directly interact with the exposed histone octamer to stabilize Cbf1-nucleosome binding. By engineering Cbf1-Pho4 chimeras, we find that while the N-terminus of the bHLH region directly contacts DNA mediating binding specificity, the C-terminal HLH region controls binding kinetics. Finally, measurements in live cells show that the Cbf1 HLH region enables efficient NDR generation by facilitating nucleosome invasion at modest TF concentrations or at lower affinity sites. Our findings provide new molecular insights into how PFs are distinct from canonical TFs so that they can efficiently invade and then displace nucleosomes.

## RESULTS

### Cbf1 pioneer activity is contained within the structured bHLHZ domain

Given our previous finding that nucleosomes slow Cbf1 dissociation (Donovan et al., 2019b), we sought to localize the region(s) responsible for nucleosome specific interactions. Since Cbf1’s unstructured N-terminus comprises approximately 60% of the protein, we first probed the impact of removing the N-terminus (Cbf1ΔN) on nucleosome binding relative to DNA **(Figure 1A, S1A).** DNA and nucleosomal templates were prepared with the consensus Cbf1 motif, where nucleosomes contain the binding site 8 bp from the nucleosome edge consistent with previous studies (**Figure S1B)** (Gibson et al., 2016; Li and Widom, 2004; Li et al., 2005). Electrophoresis mobility shift assays (EMSAs) revealed that Cbf1ΔN (residues 209-351) binds both DNA and nucleosomes over very similar concentration ranges (**Figure S1C**) as we previously observed for full length Cbf1. Furthermore, the overall affinities of Cbf1ΔN are similar to full length Cbf1 (Donovan et al., 2019b). These results indicate that, similar to our earlier characterization of the pioneer factor Reb1, the intrinsically disordered N-terminal region of Cbf1 does not contribute to overall binding affinity to DNA and nucleosomes.

**Figure 1.**
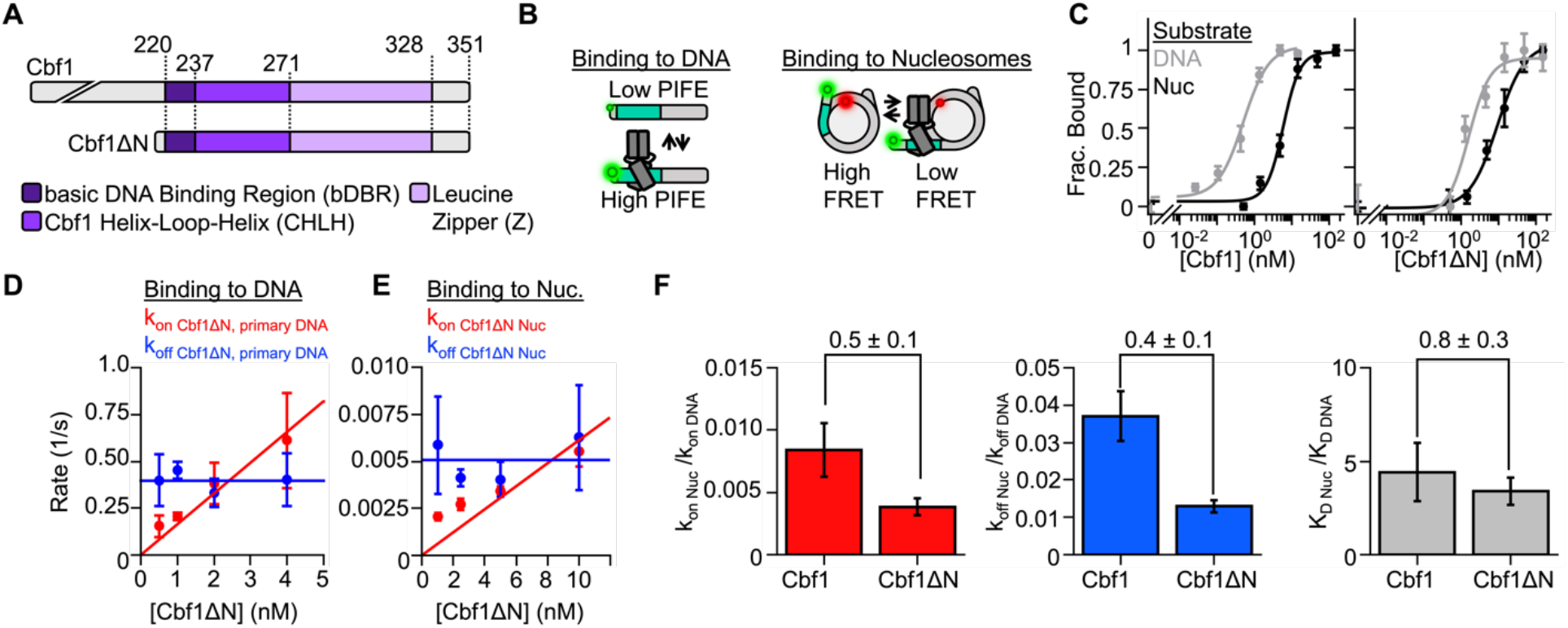
The pioneering ability of Cbf1 to efficiently invade a nucleosome is contained within the bHLHZ domain. (**A**) Domain diagrams of Cbf1 and Cbf1ΔN (AA 209-351). (**B)** Fluorescence approaches to characterize TF binding. TF-DNA interactions are detected by PIFE (Protein Induced Fluorescence Enhancement) where TF association to its target site immediately neighboring a Cy3 Fluorophore results in increased Cy3 fluorescence. TF-nucleosome interactions are detected with FRET (Fluorescence Resonance Energy Transfer) between Cy3-DNA and Cy5-H2A(K119C). FRET efficiency decreases upon TF binding to its target site within a partially unwrapped nucleosome. (**C**) Ensemble fluorescence measurements of Cbf1 (left) and Cbf1ΔN (right) binding to DNA (grey) or nucleosomes (black). Binding to DNA was detected by monitoring PIFE and binding to nucleosomes was detected by monitoring FRET efficiency between Cy3-DNA and Cy5-H2A(K119C). The PIFE and FRET data were fit to binding isotherms to determine the S_1/2_ values, which are listed in **Table S1**. (**D**) The primary Cbf1ΔN binding and dissociation rates to and from DNA at 4 Cbf1 concentrations measured by smPIFE. The dissociation rates (blue) are constant with an average rate of k_off Cbf1ΔN DNA primary_ = 0.39 ± 0.02 s^-1.^ The binding rate constant (red) is determined from the slope of a linear fit to the binding rate at four concentrations, k_on Cbf1ΔN DNA primary_ = 0.16 ± 0.01 s^-1^ nM^-1^. (**E**) Cbf1ΔN binding (red) and dissociation (blue) kinetics to and from nucleosomes, measured by smFRET. The dissociation rate (k_off Cbf1ΔN Nuc_ = 0.0051 ± 0.0006 s^-1^) and the binding rate constant (k_on Cbf1ΔN Nuc_ = 0.0006 ± 0.0001 s^-1^ nM^-1^) were determined as described for DNA. (**G**) Summary of how nucleosomes relative to DNA influence the Cbf1 and Cbf1ΔN binding rates (k_on Nuc_/k_on DNA_, red), dissociation rate constants (k_off Nuc_/k_off DNA_, blue), and binding affinity (K_D Nuc_/K_D DNA_, gray).

We next measured Cbf1ΔN binding to both DNA and nucleosomes using fluorescence-based assays which, in contrast to EMSAs, are performed in equilibrium conditions, and directly amenable to single-molecule measurements. To probe Cbf1ΔN binding to DNA, we utilized Protein Induced Fluorescence Enhancement (PIFE) (**Figure 1B, left**). In this assay, a Cy3 fluorophore is located immediately adjacent to the Cbf1 binding site so that upon Cbf1 binding, Cy3 fluorescence increases by about 1.3-fold (Hwang and Myong, 2014; Hwang et al., 2011; Luo et al., 2007). To detect nucleosome binding (**Figure 1B, right)**, we inserted a fluorophore pair between Cy3-labeled DNA and Cy5-labeled H2A(K119C). With this labeling approach, Cy3-Cy5 will undergo efficient (∼80-90%) Förster Resonance Energy Transfer (FRET) when the nucleosome is fully wrapped and transition to a lower FRET efficiency upon nucleosome partial unwrapping that is trapped by TF binding. With these approaches, we measured the concentration of Cbf1ΔN that binds 50% of DNA molecules or nucleosomes (S_1/2_). The nucleosome S_1/2_ relative to DNA is only 5-fold higher (**Table S1**, S_1/2 Cbf1ΔN DNA_ = 1.8 ± 0.6 nM, S_1/2 Cbf1ΔN Nuc_ = 9.3 ± 1.3 nM), which is similar to that of full length Cbf1 (**Figure 1C**). Both the gel- and fluorescence-based affinity measurements are in close agreement and indicate that removal of Cbf1’s N-terminus has little impact on (a) binding affinity to DNA or nucleosomes and (b) the ability to target nucleosomes via the site exposure mechanism.

Since equilibrium binding affinities are the ratio of the binding and dissociation rates, and a defining property of Cbf1 is the slow dissociation from nucleosomes, we investigated if removal of the intrinsically disordered N-terminal domain influenced the binding and dissociation kinetics. We performed single-molecule fluorescence experiments to determine Cbf1ΔN binding and dissociation kinetics to and from both DNA and nucleosomes. The kinetics to DNA were performed in a single-molecule (sm) PIFE assay where Cy3-labeled DNA was tethered to the microscope surface through a biotin-streptavidin linkage and excited via Total Internal Reflection Fluorescence (TIRF) microscopy, where high and low Cy3 fluorescence intensity represents the bound and unbound states of Cbf1ΔN, respectively (Hwang and Myong, 2014; Luo et al., 2014a). We carried out this measurement at four Cbf1ΔN concentrations and fitted raw time traces with a 2-state Hidden Markov Model (HMM) (Bronson et al., 2009) to determine the corresponding time-scales of binding (**Figure S2**). Likelihood ratio tests (Woody et al., 2016) indicate the dwell time distributions are best fit to a model that includes two characteristic dwell times in both the unbound and bound states. We focused on the faster population which comprises 70% of events (**Figure S3A-C)**.

As in previous studies (Luo et al., 2014b), we measured Cbf1ΔN binding to nucleosomes through single molecule (sm) FRET, where high and low FRET efficiency correspond to the unbound and bound states of Cbf1ΔN, respectively. The time traces **(Figure S2)** were analyzed in the same way as the smPIFE traces. The likelihood ratio tests indicated a single binding rate and dissociation rate (**Figure S4A-C)**. From the dwell time distributions **(Figure S3A, S4A)** we extracted the binding and dissociation rates of Cbf1ΔN to and from DNA and nucleosomes **(Figure 1D-E, Table S2-S3**).

As expected, the Cbf1ΔN dissociation rates from both DNA and nucleosomes did not depend on TF concentration and are fit to horizontal lines where the y-intercept represents the dissociation rate **(Figure 1D-E, blue line)**. Conversely, the binding rate increases with TF concentration, and is fit to a line where the slope represents the binding rate constant **(Figure 1D-E, red line)**. Importantly, both the binding and dissociation rates of Cbf1ΔN to and from nucleosomes relative to DNA are within a factor of 2 compared to full length Cbf1 (**Figure 1F**, **Table S4**). These results imply that the structured bHLHZ domain of Cbf1, but not the disordered N-terminal domain, is responsible for strong nucleosome binding.

### The basic helix-loop-helix structural motif can invade nucleosomes as either a pioneer or canonical transcription factor

To further understand the mechanism behind the slow dissociation of Cbf1 from nucleosomes, we investigated a second bHLH TF from budding yeast, Pho4, to provide a distinct comparison. Pho4 also recognizes an E-box motif via the N-terminal region of the bHLH domain (**Figure 2A)** (Cave et al., 2000; Maerkl and Quake, 2007; Shimizu et al., 1997) and, unlike Cbf1, Pho4 occupancy is strongly inhibited by nucleosomes *in vivo* (Rossi et al., 2018; Zhou and O’Shea, 2011). To provide a quantitative comparison, we first investigated the affinity of Pho4 to its consensus motif on DNA with EMSAs. Unless otherwise noted, all subsequent experiments matched the Cbf1 or Pho4 consensus target sequence with the corresponding Cbf1 or Pho4 basic region. In this assay, we found that the DNA S_1/2_ is approximately equal to the total DNA concentration of 0.2 nM **(Figure S5A)**. This implies that Pho4 binds DNA stoichiometrically and that the S_1/2_ does not accurately reflect the dissociation constant (K_D_). Instead, here the S_1/2_ is an upper limit of the apparent K_D_ for DNA binding (Jarmoskaite et al., 2020). We then investigated the Pho4 binding affinity to DNA with ensemble PIFE, which enables experiments at a lower DNA concentration of 50 pM of Cy3 labeled DNA. Even at this lower DNA concentration, we measured a Pho4 binding S_1/2_ that is within the uncertainty of the total DNA concentration (S_1/2 Pho4 DNA_ = 30 ± 20 pM, **Figure 2B, left).** Therefore, we conclude that the apparent K_D_ of Pho4 binding to naked DNA is significantly below 30 pM, reflecting a very high binding affinity.

**Figure 2:**
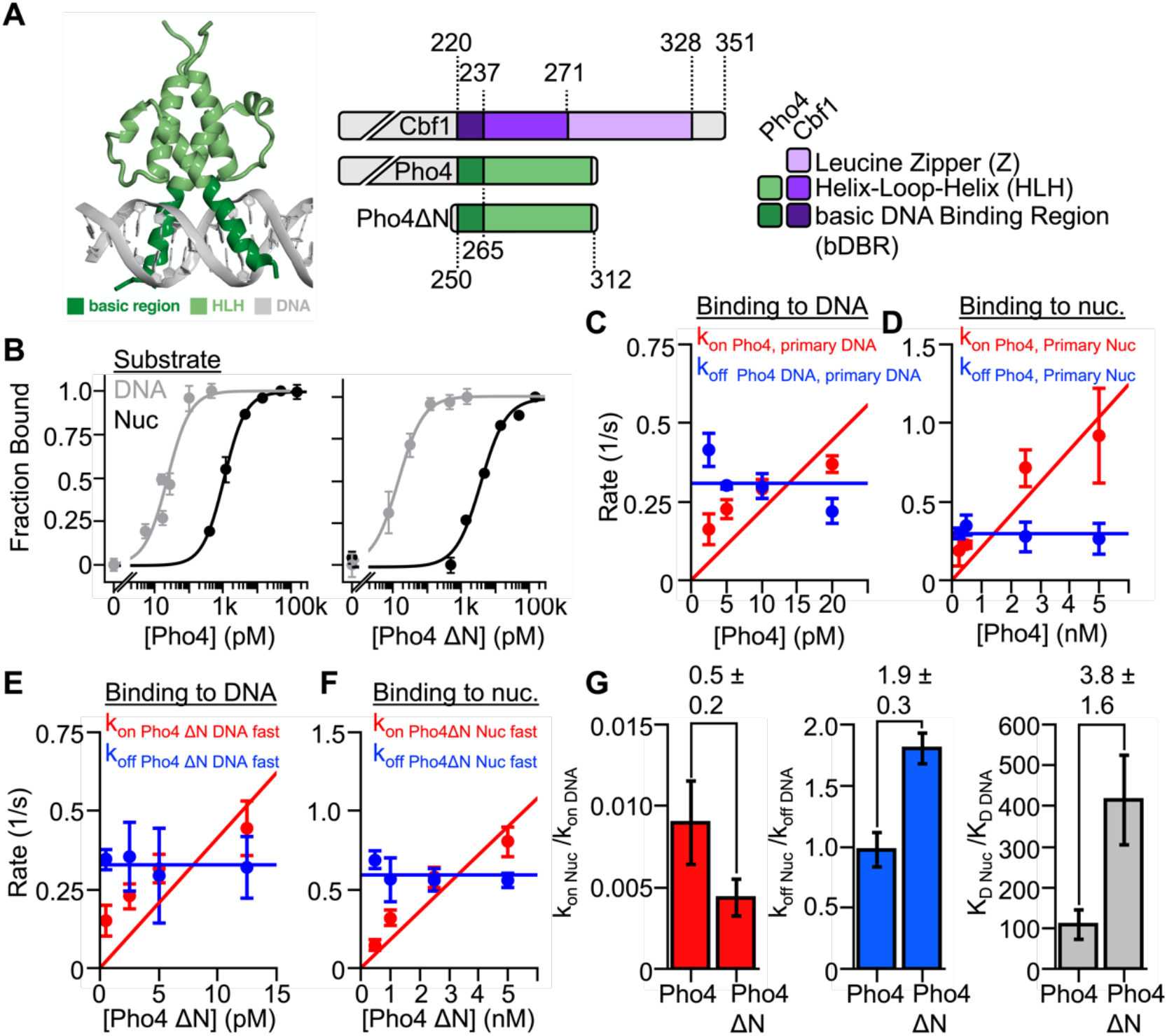
Nucleosomes restrict Pho4 binding by more than 100-fold. (**A**) Crystal structure of bHLH TF Pho4 (Shimizu et al., 1997) and domain diagrams for Cbf1, Pho4, and Pho4ΔN. (**B**) Ensemble PIFE (grey) and FRET (black) measurements of full length Pho4 (left) and Pho4ΔN (right) binding their consensus target sequence within DNA and nucleosomes. Binding isotherm fits determine the S_1/2_ of each titration (S_1/2 Pho4 DNA_ = 30 ± 20 pM, S_1/2 Pho4ΔN DNA_ = 15 ± 1 pM, S_1/2 Pho4 Nuc_ = 1.08 ± 0.04 nM, S_1/2 Pho4ΔN Nuc_ = 3.88 ± 0.93 nM). (**C**) Summary of the smPIFE studies of full length Pho4 binding and dissociation kinetics to and from DNA, which are plotted and analyzed in the same way as Figure 1D (k_on Pho4 DNA primary_ = 20 ± 5 s^-1^ nM^-1^, koff _Pho4 DNA primary_ = 0.31 ± 0.04 s^-1^). (**D**) Summary of the smFRET studies of Pho4 binding and dissociation kinetics to and from nucleosomes, plotted and analyzed in the same way as Figure 1E (k_on Pho4 Nuc primary_ = 0.20 ± 0.04 s^-1^ nM^-1^, k_off Pho4 Nuc primary_ = 0.30 ± 0.02 s^-1^. (**E**) Summary of the smPIFE studies of Pho4ΔN binding and dissociation kinetics to and from DNA, which are plotted and analyzed in the same way as figure 1D (k_on Pho4ΔN DNA primary_= 40 ± 10 s^-1^ nM^-1^, k_off Pho4ΔN DNA primary_ = 0.33 ± 0.01 s^-1^. (**F**) Summary of the smFRET studies of Pho4 binding and dissociation kinetics to and from nucleosomes, plotted and analyzed in the same way as Figure 1E (k_on Pho4ΔN Nuc_ = 0.18 ± 0.02 s^-1^ nM^-1^, k_off Pho4ΔN Nuc_ = 0.59 ± 0.03 s^-1^). (**G**) Summary of the nucleosome binding kinetics (red), dissociation kinetics (blue) and dissociation constant (gray) relative to DNA for both full length Pho4 and Pho4ΔN.

We next characterized Pho4 binding to its consensus sequence within a nucleosome using both EMSAs (**Figure S2B, bottom**) and ensemble FRET (**Figure 2B, left**). In contrast to Pho4-DNA affinity measurements, Pho4-nucleosome affinity measurements occur over a sufficiently high concentration range that the S_1/2_ accurately represents the apparent K_D_. Monitoring FRET while titrating Pho4 reveals Pho4 binds via the site exposure mechanism with an S_1/2_ consistent with our EMSA measurement (S_1/2 Pho4 nuc_ = 1.08 ± 0.04 nM) (**Figure 2B left panel)** indicating that nucleosomes impede Pho4 binding by much more than 40-fold (**Table S1**). We conclude that despite both Cbf1 and Pho4 being bHLH TFs that bind nucleosomes via the site exposure model, Cbf1 efficiently binds within nucleosomes while Pho4 binding is strongly inhibited by nucleosomes.

To understand the difference in Cbf1 and Pho4 nucleosome affinity, we considered the possibility that the bHLH region of Pho4 binds nucleosomes with similar efficiency as Cbf1ΔN, but that the disordered N-terminal domain of Pho4 inhibits efficient binding. To test this, we purified Pho4ΔN (AA 250-312) and measured its binding affinity towards DNA and nucleosomes. Our results agree with previous crystallographic studies indicating this mutation does not greatly impact DNA binding (Shimizu et al., 1997) as we still measure stoichiometric binding to DNA **(Figure 2B, right; Figure S5C)**. Furthermore, Pho4ΔN binds nucleosomes even weaker than the full length Pho4 (S_1/2 Pho4 ΔN Nuc_ = 3.88 ± 0.93 nM) (**Figure 2B, S5D**). These results indicate that, similar to Cbf1, the unstructured N-terminus plays little role in the differential affinity of Pho4 to nucleosomes and DNA. Furthermore, this shows that the difference in nucleosome binding efficiency between Cbf1 and Pho4 are contained within their bHLH regions.

Reb1 and Cbf1 efficiently bind to nucleosomes because of their slow dissociation rate relative to DNA (Donovan et al., 2019b), while the canonical TF Gal4 binding within nucleosomes is strongly inhibited because it is accelerated off its target site by the nucleosome (Luo et al., 2014a). We therefore investigated the possibility that the difference in binding affinities to nucleosomes between Cbf1 and Pho4 is also due to differences in dissociation rates from nucleosomes. We performed smPIFE and smFRET measurements to determine full-length Pho4 and Pho4ΔN binding and dissociation rates to and from its target site within DNA and nucleosomes (**Figures S6-S9**). The cumulative sums of Pho4 binding and dissociation kinetics to and from DNA and nucleosomes are best fit by two exponentials (**Figure S6-S8**). In addition, the cumulative sums of Pho4ΔN kinetics to and from DNA are best fit by two exponentials (**Figure S8**), while the kinetics to and from nucleosomes are best fit by a single exponential (**Figure S9**). The primary rates are summarized in **Tables S2** and **S3**. We find that the nucleosome reduces the binding rate constant of both full-length Pho4 and Pho4ΔN by about 2 orders of magnitude relative to DNA (**Figure 2C-G)**. This is nearly identical to other TFs binding rates to the same location within the nucleosome relative to DNA including Cbf1, Reb1 and LexA (Donovan et al., 2019b; Luo et al., 2014a), and is consistent with previous genome-wide ChIP-seq experiments indicating nucleosomes drastically restrict Pho4 binding (Zhou and O’Shea, 2011). However, in contrast to Cbf1 the dissociation rate of full-length Pho4 is the same from nucleosomes and DNA, while Pho4ΔN is *accelerated* off nucleosomes relative to DNA by a factor of two **(Figure 2G)**. Overall, these results show that differences between the bHLH domains of Cbf1 and Pho4 result in these TFs having very different interactions with nucleosomes. Cbf1 efficiently invades the nucleosome via the dissociation rate compensation mechanism, while Pho4 occupancy is strongly suppressed by the nucleosome.

### The cryo-EM structure of Cbf1 bound to its site within the nucleosome reveals specific interactions between the Cbf1 HLH region and the histone octamer

We hypothesized the slowed dissociation from nucleosomes by Cbf1 is accomplished through additional interactions with the nucleosome compared to DNA. To test this hypothesis, we resolved the structure of the Cbf1-nucleosome complex using cryo electron microscopy (cryo-EM) single particle analysis (**Figure 3A-B**). We used full-length Cbf1 protein together with the same consensus Cbf1 motif engineered at the same position within the Widom 601 nucleosome DNA positioning sequence as the previous in vitro binding experiments. The Cbf1-nucleosome complex was reconstituted, purified by size exclusion chromatography and vitrified without crosslinking. 6339 movies collected on a 300 keV Krios microscope were processed to yield 1.7M particles of the complex. Due to considerable heterogeneity in the Cbf1 region of the map, we employed 3D classification to obtain a more homogenous subset of particles. Reconstructions of subsets containing several hundred thousand particles reached 2.7 Å overall resolution, limited by the Nyquist frequency of the binned images. The data showed that the volume of the nucleosome core was well-resolved but the region corresponding to Cbf1 displayed considerable flexibility. Multiple rounds of template-guided 3D classification in cryoSPARC, and local classification without alignment in Relion were performed to obtain a more homogenous subset of particles. The final subset that was used to build the model consisted of 73k particles and yielded a map with 3.2-Å overall resolution and a local resolution of 4–7 Å in the Cbf1 region (**Figure 3C**).

**Figure 3.**
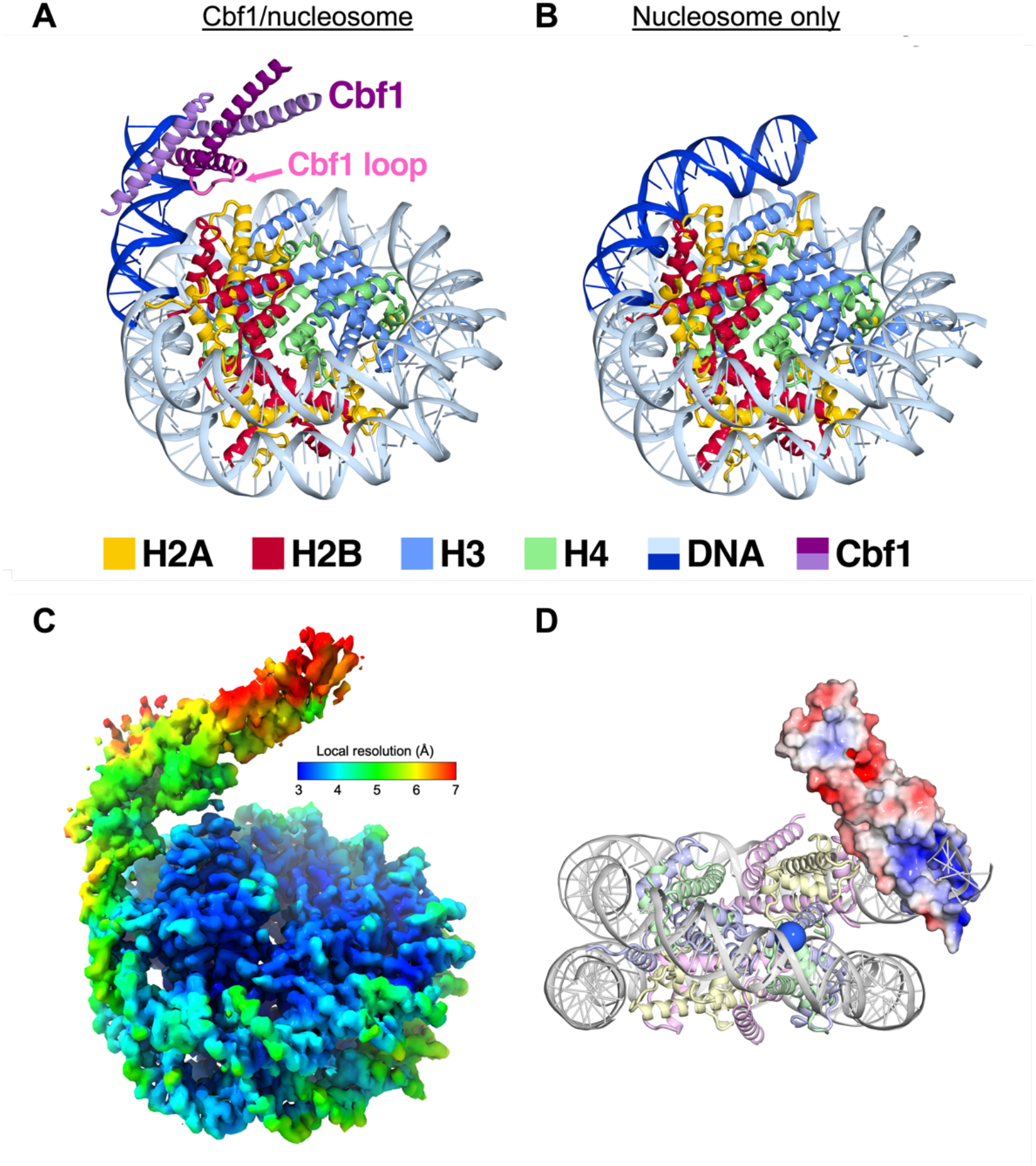
Cryo-EM structure of the Cbf1-nucleosome complex reveals specific interactions between the Cbf1 HLH domain and histones. (A) Ribbon representation of Cbf1-nucleosome complex at 5-Å resolution. The Cbf1 dimer is colored in light and dark purple. The 20 bp of unwrapped DNA is in dark blue, while the DNA in contact with the histone octamer is colored in sky blue. Histone H2A, H2B, H3 and H4 are colored in yellow, red, light blue and light green, respectively. (B) A ribbon representation of the fully wrapped nucleosome (Vasudevan et al., 2010). The coloring is the same as in panel (A). (C) The 3.2-Å 3D reconstruction used for model building colored by local resolution, calculated with cryoSPARC. (D) Cbf1 helix 2 displays negatively charged patches (red) that could interact with positively charged histone tails. The APBS-calculated electrostatic (Jurrus et al., 2018) potential was mapped from −5 to 5 kT/e. The view faces the nucleosome dyad. The Cα of the first modeled histone H3 residue, Pro43, is rendered as a blue sphere.

The data supports a model where the homodimer of the basic helix-loop-helix Leu-zipper (bHLHZ) domain of Cbf1 (residues 222-288) binds the E-box motif near the nucleosome entry-exit site. In agreement with our FRET studies, our model shows how Cbf1 traps the nucleosome in an unwrapped state with at least 20 bp of DNA peeled off the histone octamer (**Figure 3A** vs. **Figure 3B**) (Vasudevan et al., 2010). In addition to unwrapping, the nucleosomal DNA occupied by Cbf1 is angled outward, away from the histone core. In this conformation, the Cbf1 HLH region is positioned to directly interface with the octamer. Extensive 3D classification yielded particle classes where Cbf1 and the unwrapped DNA are positioned at various angles relative to the nucleosome core (**Figure S10A**). The continuous nature of the structural heterogeneity does not allow further improvement of the map portion corresponding to Cbf1. We were able to reach a local resolution of about 4 Å in the vicinity of the interacting histones, but the resolution drops off to 5 Å and worse in the periphery (**Figure 3C**).

Despite the dynamic nature of this structure, which makes it difficult to resolve Cbf1-histone contacts, this model articulates the important regions for interaction. The loop of the HLH domain appears to be positioned closest to the core histones. In particular, we observe map density corresponding to an interaction between Cbf1 E253 and H2A K75 on the H2A histone fold L2 loop, indicative of one of potentially several interactions between Cbf1 and core histones (**Figure S10B and S10C**). In fact, the entire HLH region features many negatively charged solvent exposed amino acids capable of electrostatically interacting with the histone core and/or histone tails (**Figure 3D**). These electrostatic interactions between Cbf1 and histones in combination with Cbf1-DNA binding likely results in the observed motion of Cbf1 relative to the nucleosome (**Figure S10A**). This cluster of interactions is the fulcrum point about which Cbf1 and the unwrapped DNA pivot, acting like a ball joint. Overall, these results provide direct evidence that Cbf1 interacts with the histone octamer and provides a mechanism for its slow dissociation from the nucleosome.

### The addition of Cbf1 Leucine Zipper to Pho4 does not enhance DNA or nucleosome binding

Our single-molecule measurements indicate that only the structured regions comprising the bHLHZ and bHLH regions for Cbf1 and Pho4, respectively, are important for binding DNA and nucleosomes. However, much of the Cbf1 leucine zipper is unresolved in our Cryo-EM structure. We therefore considered the possibility that the C-terminal leucine zipper extension of Cbf1, which Pho4 does not contain, is important for efficient binding within nucleosomes. To test this hypothesis, we fused Cbf1 residues 270-351 onto Pho4’s C-terminus (Pho4 Zip, **Figure S11A)**. Just like full-length Pho4, we measured Pho4 Zip binding to the Pho4 consensus motif within DNA and nucleosomes. We observe stoichiometric binding of Pho4 Zip to its target site within DNA (**Figure S11B**), while Pho4 Zip binds nucleosomes with the same affinity as full-length Pho4 (**Figure S11C**). This indicates that the Cbf1 leucine zipper, while important for dimerization, does not contribute to the increased efficiency of binding within nucleosomes. This result is consistent with the cryo-EM structure where the start of the leucine zipper is positioned so that it extends away from the core histones.

### The HLH region regulates the dissociation rate from nucleosomes

Our structure of the Cbf1-nucleosome complex indicates the HLH region interacts directly with the histone octamer. To further investigate the role of this region in controlling binding kinetics, we attempted to eliminate the HLH-nucleosome interaction. To do this, we generated a Cbf1-Pho4 chimera where the Cbf1 HLH region, residues 237-351, is replaced with the corresponding region in Pho4, residues 265-312 **(Figure 4A)**. We refer to this chimera as Cbf1ΔN PHLH, where “PHLH” represents the “Pho4 Helix-Loop-Helix” dimerization region. Using ensemble PIFE measurements, we find that Cbf1ΔN PHLH binds stoichiometrically to the Cbf1 consensus motif on naked DNA with a S_1/2_ of 37 ± 5 pM (**Figure S12A-B**). We then used ensemble FRET measurements to determine that the S_1/2_ for Cbf1ΔN PHLH binding to nucleosomes is 3.1 ± 0.5 nM (**Figure S12C**, **Table S1)**, representing an 80 ± 10-fold increase from that of DNA (**Table S1**). In comparison, Cbf1ΔN binding to nucleosomes relative to DNA (S_1/2 Nuc_ / S_1/2 DNA_) is only 3.4 ± 0.7-fold higher. These results show that, in contrast to Cbf1ΔN, Cbf1ΔN PHLH chimera behaves more like Pho4ΔN, where it binds tightly to the Cbf1 motif but is unable to efficiently invade into the nucleosome.

**Figure 4:**
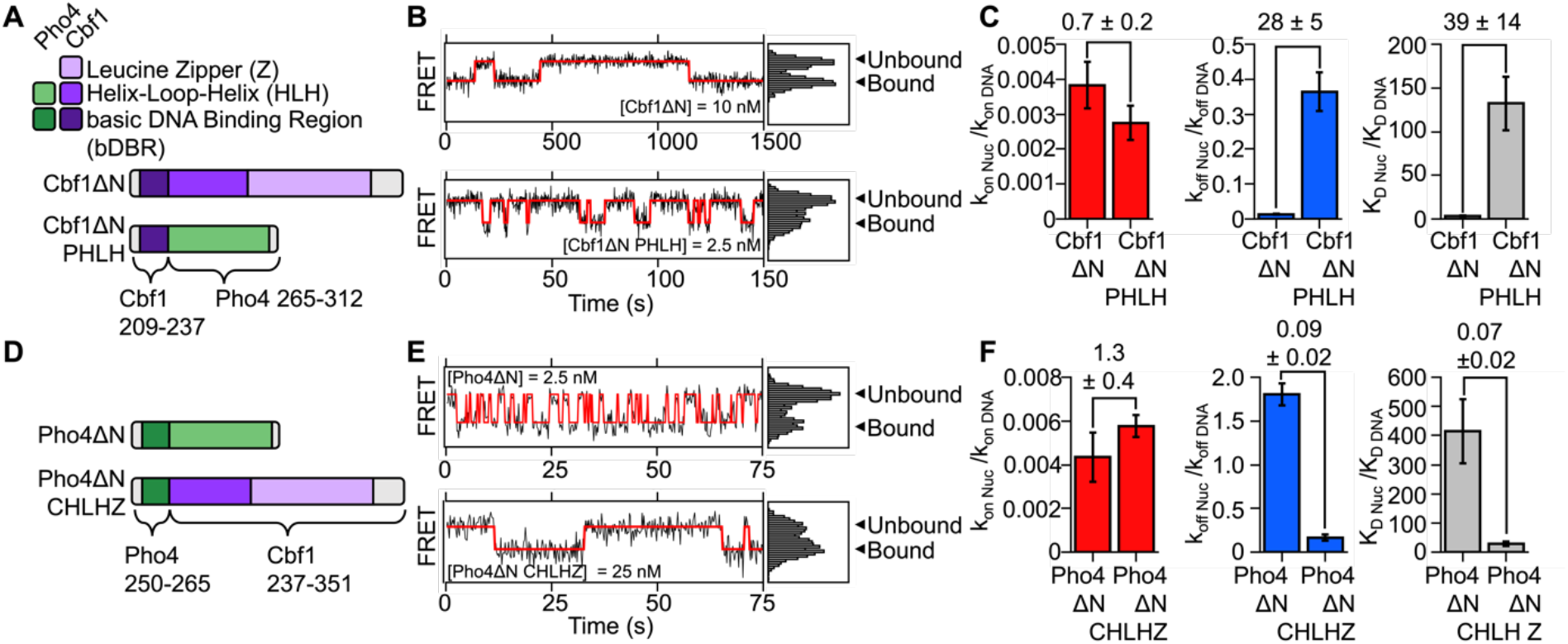
The Cbf1 HLH region facilitates pioneer activity through the dissociation rate compensation mechanism. (**A**) Domain diagram of Cbf1ΔN and the Cbf1ΔN PHLH chimera where Cbf1 residues 237-351 was swapped out for Pho4 residues 264-312. (**B**) Representative smFRET time traces of Cbf1ΔN (top) and Cbf1ΔN PHLH (bottom) binding to and dissociating from nucleosomes. Note that the Cbf1ΔN time axis is 10 times longer than the Cbf1ΔN PHLH time axis. Visual comparison reveals the drastically slower dissociation rate of Cbf1ΔN relative to Cbf1ΔN PHLH. (**C**) Comparison of nucleosome binding rates (red), dissociation rates (blue) and dissociation rates relative to DNA for Cbf1ΔN and Cbf1ΔN PHLH. (**D**) Domain diagram of Pho4ΔN and the Pho4ΔN CHLHZ chimera where Pho4 265-312 was swapped out for Cbf1 residues 237-351. (**E**) Representative smFRET time traces of Pho4ΔN (top) and Pho4ΔN CHLHZ (bottom) binding to and dissociating from nucleosomes. Visual inspection of these time traces reveals the drastically faster dissociation rate of Pho4ΔN relative to Pho4ΔN CHLHZ. (**F**) Comparison of nucleosome binding rates (red), dissociation rates (blue) and dissociation rates relative to DNA for Pho4ΔN and Pho4ΔN CHLHZ.

We then investigated if the low relative affinity of Cbf1ΔN PHLH towards the nucleosome results from the loss of the dissociation rate compensation mechanism. SmPIFE experiments were used to determine the Cbf1ΔN PHLH binding rate constant and dissociation rate to and from the Cbf1 consensus motif within DNA (**Figure S13, Table S2**). Separately, smFRET measurements were used to determine the Cbf1ΔN PHLH binding rate constant and dissociation rate to and from nucleosomes (**Figure S14, Table S3**). Visual inspection of time traces shows many binding fluctuations per nucleosome during one video, indicating that Cbf1ΔN PHLH bound dwell times are much shorter to nucleosomes compared to the long dwell times characteristic of Cbf1ΔN (**Figure 4B, S2A vs S2D**). Quantification of the measured dwell times reveals that nucleosomes reduce the binding rate of Cbf1ΔN PHLH by 360-fold (k_on nuc_ / k_on DNA_ = 0.0027 ± 0.0005), which is similar to the reduction with Cbf1ΔN **(Figure 4C, S13-S14, Table S3)**. However, the Cbf1ΔN PHLH dissociation rate is only 3-fold slower from nucleosomes relative to DNA (k_off nuc_ / k_off DNA_ = 0.36 ± 0.05), while the Cbf1ΔN dissociation rate is 80-fold slower from nucleosomes relative to DNA (k_off nuc_ / k_off DNA_ = 0.013 ± 0.002, **Figure 4C**, **Table S3**). These results show that the replacement of the Cbf1 HLH region with the Pho4 HLH region results in a similar dissociation rate from nucleosomes as compared to DNA and that this loss in the dissociation rate compensation results in nucleosomes more strongly inhibiting binding of Cbf1ΔN PHLH.

Our studies of Cbf1ΔN PHLH imply that Cbf1 residues 237-351, in addition to mediating dimerization, are responsible for slowed dissociation from nucleosomes. To further investigate this idea, we prepared another Cbf1-Pho4 chimera, Pho4ΔN CHLHZ (CHLHZ = Cbf1 Helix-Loop-Helix dimerization region), where the Pho4 HLH region (residues 265-312) is replaced with the Cbf1 HLH region (residues 237-351) (**Figure 4D, S15A**). The combination of ensemble PIFE measurements of DNA binding and FRET measurements of nucleosome binding reveal that nucleosomes reduce Pho4ΔN CHLHZ binding by 40-fold (S_1/2 Nuc_ / S_1/2 DNA_ = 40 ± 20), which is a 7-fold reduction in relative nucleosome binding affinity compared to Pho4ΔN (S_1/2 Nuc_ / S_1/2 DNA_ > 270 ± 70) (**Figure S15, Table S1**). These results indicate that addition of the Cbf1 HLH region facilitates nucleosome binding.

To directly probe if Pho4ΔN CHLHZ acquired efficient nucleosome binding within the nucleosome via the dissociation rate compensation mechanism, we performed single-molecule measurements of Pho4ΔN CHLHZ binding and dissociation dynamics to and from DNA **(Figure S16, Table S2)** and nucleosomes **(Figure S17, Table S3)** as we did with Cbf1ΔN PHLH. We found that the Pho4ΔN CHLHZ binding rate to its site within the nucleosome is reduced by 170-fold relative to the DNA binding rate (k_on nuc_ / k_on DNA_ = 0.0058 ± 0.0005). This is similar to the 250-fold reduction in nucleosome binding rate relative to DNA for Pho4ΔN. In contrast, unlike Pho4ΔN, which dissociates 2-fold *faster* from nucleosomes (k_off nuc_ / k_off DNA_ = 1.8 ± 0.1), the dissociation rate of Pho4ΔN CHLHZ from nucleosomes is *slower* by 6-fold relative to the dissociation rate from DNA (k_off nuc_ / k_off DNA_ = 0.16 ± 0.04, **Figure 4E-F, Table S4**). This is more than a factor of 10-fold reduction in the nucleosome dissociation rate relative to DNA between Pho4ΔN CHLHZ and Pho4ΔN. Taken together, these results show that the HLH dimerization region of Cbf1 enables the slow nucleosome dissociation rate of Pho4ΔN CHLHZ resulting in efficient binding within the nucleosome via the dissociation rate compensation mechanism. These combined studies of Cbf1-Pho4 chimeras suggest that the HLH region of Cbf1 provides additional interactions with the nucleosome, which that are absent in Pho4 HLH region, and contribute to efficient targeting of Cbf1 to its binding site within the entry-exit region of the nucleosome.

### The Cbf1 HLH region imparts nucleosome displacement *in vivo*

To address how slowed dissociation from nucleosomes imparted by the Cbf1 HLH region relates to function in live cells, we probed the ability of these chimeric TFs to establish nucleosome depleted regions (NDRs) by generating yeast strains with a few genetic modifications **(Table S5, Figure S18)**. First, we introduced Pho4ΔN, Cbf1ΔN, or chimeras driven by the *MET3* promoter into yeast so that these factors can be induced by the absence of methionine. In addition to a nuclear localization signal (NLS), these TFs contain a GFP on their C-terminal to validate protein expression and localization. Note that Pho4ΔN lacks the phosphorylation sites in the full-length Pho4 that are required for their cytoplasmic localization in high phosphate media (O’Neill et al., 1996). Each of these factors were nuclear-localized and expressed to similar levels upon methionine depletion (**Figure S18A**). Second, to eliminate the effect from endogenous Cbf1, we replaced the *CBF1* promoter with the *Gal1* promoter so that the wt Cbf1 is effectively depleted in glucose media. The endogenous Pho4 was intact in our strains, but all the following measurements were performed in high phosphate conditions where Pho4 was sequestered in the cytoplasm (O’Neill et al., 1996). Finally, on a mutated *HO* promoter assembled into a constitutive nucleosome array (nucleosome −1 to −7) (Yan et al., 2018), we engineered a single Pho4 or Cbf1 binding site 43 bp from the dyad into nucleosome - 4. In this way, accessing these motifs by Pho4 or Cbf1 variants requires direct competition with the nucleosome.

Using these strains, we assayed how these mutant TFs influence nucleosome positioning near the engineered motifs using an MNase digestion assay followed by stacking qPCR. On the template containing a Cbf1 motif, we observed a clear shift in nucleosome positioning and the formation of a short NDR over the Cbf1 motif upon expression of Cbf1ΔN, indicating that Cbf1ΔN, like the full-length Cbf1, has nucleosome displacement activity *in vivo* (**Figure 5A**). No discernible changes in nucleosome positioning were observed upon induction of Pho4ΔN **(Figure 5A, panel 2/4)**. Remarkably, swapping the HLH region of the two factors reversed their nucleosome invasion activity: Pho4ΔN CHLHZ, but not Cbf1ΔN PHLH, was able to shift nucleosome positioning near the Cbf1 motif **(Figure 5A, bottom two panels)**. These results are highly correlated with our *in vitro* observations that Pho4ΔN CHLHZ has a slower dissociation rate from nucleosomes than Pho4ΔN, whereas Cbf1ΔN PHLH dissociates faster than Cbf1ΔN.

**Figure 5.**
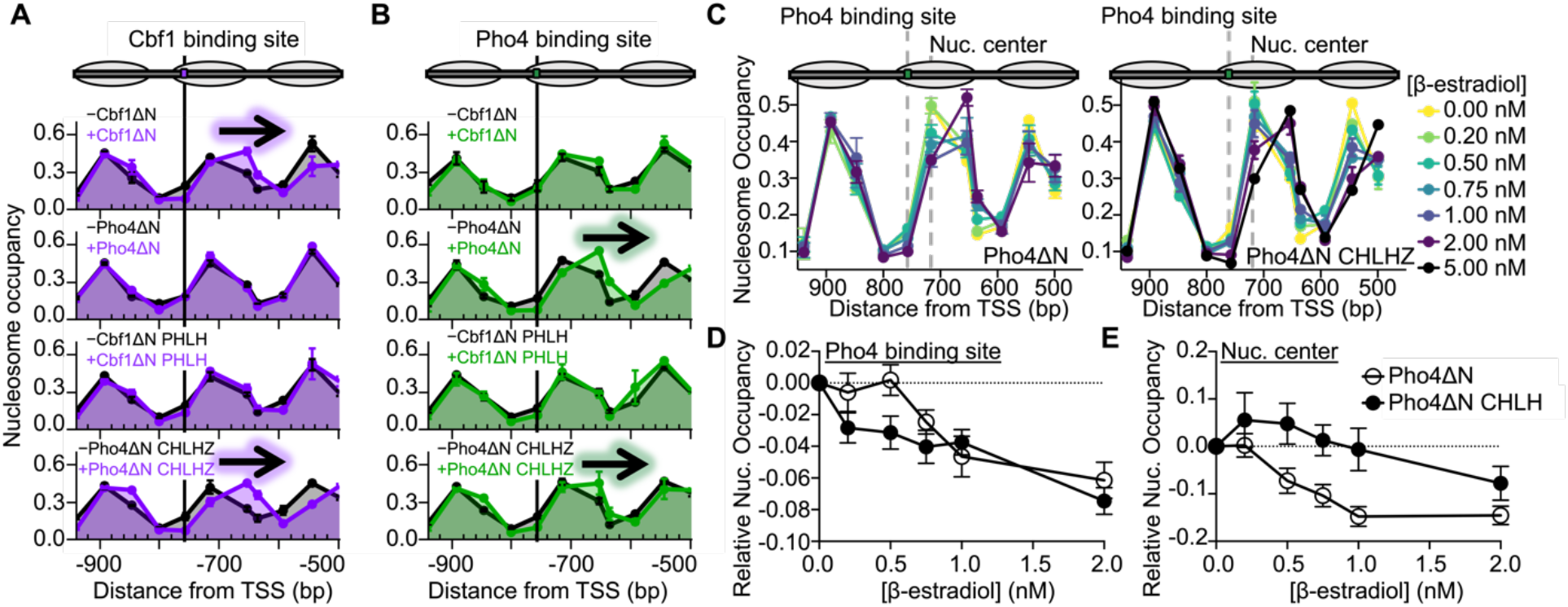
The Cbf1 HLH region imparts efficient nucleosome targeting and displacement *in vivo*. (**A**) and (**B**) Nucleosome occupancy within the *HO* promoter that contains a Cbf1 or Pho4 target site, respectively. The target sites were inserted 43 bp upstream of the −4 nucleosome dyad. The position of the Cbf1 and Pho4 target sites are indicated by the black line in (A) and (B), respectively. Nucleosome occupancy was determined in the presence (black) or absence of methionine (purple or green), where methionine depletion induces the expression of either Cbf1ΔN, Pho4ΔN, Cbf1ΔN PHLH, or Pho4ΔN CHLHZ. The arrow indicates which TFs induce a change in the position of the −4 nucleosome. (**C**) Nucleosome occupancy within the *HO* promoter with the Pho4 target sequence in the same location as panel (A). Different doses of β-estradiol titrate the expression level of Pho4ΔN (left) or Pho4ΔN CHLHZ (right), respectively, which are under the control of the *GAL1* promoter that is activated by Gal4DBD-ER-VP16. The grey dotted lines indicate the location of the Pho4 target sequence (-775 bp) and the nucleosome center without TF expression (-716 bp). (**D**) Relative nucleosome occupancy at the Pho4 binding site (-757 bp from the TSS) upon induction of Pho4 ΔN (white) or Pho4 ΔN CHLHZ (black) with increasing levels of β-estradiol. (**E**) Relative nucleosome occupancy measured at the center of the −4 nucleosome (-716 bp from the TSS) with increasing levels of β-estradiol.

We also assayed how these proteins influence nucleosome positioning when targeting the Pho4 binding site inside the nucleosome −4 (**Figure 5B**). Not surprisingly, Cbf1ΔN and Cbf1ΔN PHLH, which bind 300-fold weaker to the Pho4 site *in vitro* **(Figure S18B)**, are unable to invade nucleosomes containing the Pho4 binding site and have no influence on nucleosome positioning. Interestingly, we observe a shift in nucleosome positioning upon expression of Pho4ΔN (**Figure 5B, panel 2/4)**, which is surprising given that Pho4 has previously been reported to be blocked by nucleosomes (Zhou and O’Shea, 2011). However, this is consistent with other very tight DNA binders like the bacterial TF TetR that can also generate small NDRs (Yan et al., 2018). Finally, we also observe a clear shift in nucleosome positioning when expressing Pho4ΔN CHLHZ **(figure 5B, bottom panel)**. Taken together these data indicate two mechanisms by which TFs invade their binding sites within nucleosomes: (1) through very tight DNA binding as in the case of Pho4ΔN and (2) through the dissociation rate compensation mechanism as in the case of TFs possessing the Cbf1 HLH region.

The experiments above show that both Pho4ΔN and Pho4ΔN CHLHZ can displace nucleosomes when highly expressed in the cells. To further investigate the functional differences between these two factors, we compared the nucleosome displacement by Pho4ΔN and Pho4ΔN CHLHZ at lower concentrations. We constructed strains with either factor under control of a *GAL1* promoter activated by Gal4DBD-ER-VP16 (Louvion et al., 1993). This way, Pho4ΔN or Pho4ΔN CHLHZ expression level can be modulated by the concentration of β-estradiol **(Figure S18C)**. We observed concentration-dependent nucleosome displacement activities in both cases, where higher levels of these factors lead to a decrease of the nucleosome occupancy near their binding sites and a shift of the downstream nucleosome away from the binding sites (**Figure 5C**). For Pho4ΔN, these two events are well correlated, and both start to occur ∼ 0.5-1 nM of β-estradiol **(Figure 5D-E**). For Pho4ΔN CHLHZ, however, 0.2 nM of β-estradiol is sufficient to reduce nucleosome occupancy near its binding site, but the nucleosome shift requires 2 nM of β-estradiol. This observation indicates that, at low concentration, Pho4ΔN CHLHZ may bind within the nucleosome through DNA unwrapping, which increases the local DNA accessibility without nucleosome translocation. This is consistent with the more efficient Pho4ΔN CHLHZ-nucleosome binding observed *in vitro*. Higher levels of Pho4ΔN CHLHZ traps the nucleosome more in such a partially unwrapped unstable state, which may eventually lead to the recruitment of downstream factors and nucleosome movement. In contrast, Pho4ΔN has shorter dwell times on the nucleosome, and its binding appears to be more mutually exclusive with the nucleosome.

## DISCUSSION

Pioneer TFs are typically the first proteins to bind within a silenced gene to initiate transcription activation and therefore require the ability to access DNA target sites within nucleosomes to function (Iwafuchi-doi and Zaret, 2014; Soufi et al., 2014; Zaret and Carroll, 2011). In contrast, canonical TFs typically contribute to transcription activation after their DNA target site is accessible since nucleosomes significantly limit binding access to their target sequence (Polach and Widom, 1995). While the structure of a protein typically defines its function, transcription factors with the same structural motif can function very differently, i.e. pioneer vs. canonical TF.

In this study, we provide mechanistic insight into how two bHLH TFs, Cbf1 and Pho4, function as significantly different TFs: pioneer vs. canonical (**Figure 6A**). We find that Cbf1 has comparable affinities towards nucleosome and naked DNA through a dissociation rate compensation mechanism. Neither the N-terminal intrinsically disordered region, the C-terminal leucine zipper extension, nor the basic DNA binding region that directly interacts with the DNA motif contribute to the slow dissociation of Cbf1 from the nucleosome. Instead, our data reveals that the Cbf1 HLH region enables efficient nucleosome invasion via the dissociation rate compensation mechanism. In contrast, the corresponding HLH region of Pho4 does not impart slowed dissociation to facilitate nucleosome invasion. Importantly, we show that these HLH regions of Cbf1 and Pho4 can be swapped, resulting in the transfer of the nucleosome dissociation and invasion properties between TFs (**Figure 6B**). This indicates that the dimerization region of the bHLH domain is the pioneering module.

**Figure 6.**
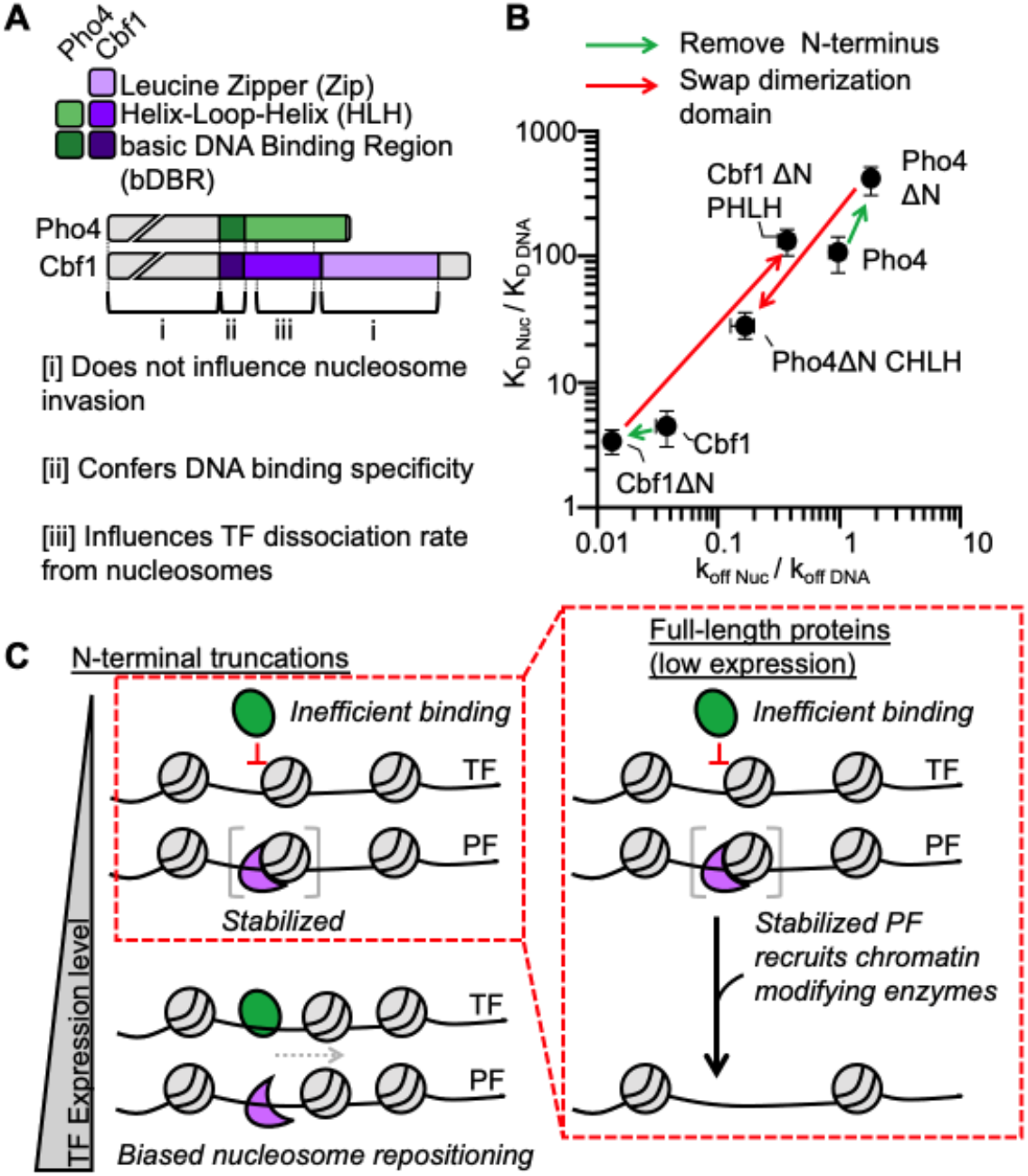
The Cbf1 and Pho4 HLH regions define their ability to invade nucleosomes *in vitro* and reposition nucleosomes *in vivo*. **(A)** Domain diagrams of full-length Pho4 and Cbf1 with the regions denoted by brackets on how they influence nucleosome invasion. (**B**) Log-log plot relating the efficiency of nucleosome invasion (K_D Nuc_/K_D DNA_, y-axis) to the relative dissociation rate (k_off Nuc_ /k_off DNA_, x-axis). Each point on this plot represents a different measurement from this study or (Donovan et al., 2019). The arrows between points indicate how specific mutations change relative dissociation rate and the corresponding efficiency of nucleosome invasion. (**C**) Model of nucleosome invasion and displacement by bHLH PFs that rely on the dissociation rate compensation mechanism and canonical bHLH TFs that are inhibited by nucleosomes. At low expression levels where binding sites are only partially occupied, bHLH PFs (purple) with their N-terminal domains deleted invade and stabilize the nucleosome, while canonical bHLH TFs (green) are blocked by the nucleosome. If the bHLH PF contains the N-terminal domain, it may recruit chromatin modifying complexes that likely generate the larger NDRs previously reported (Yan et al., 2018). However, at high expression levels both bHLH PFs and canonical bHLH TFs fully occupy their target sites within nucleosomes. If the bHLH PF or canonical bHLH TF has the N-terminal domain deleted, its occupancy biases nucleosome repositioning generating a small NDR as observed in this study.

Our three-dimensional structure of the Cbf1-nucleosome complex provides further molecular explanations for these observations. The structure confirms that the Cbf1 bHLH domain’s interactions with DNA are similar to other bHLH TFs including Pho4. The structure shows for the first time that the Cbf1 HLH region directly interacts with histones H2A and H2B, consistent with the conclusion from our single molecule chimera studies that the pioneer region of Cbf1 is contained within its HLH region. In addition, the acidic region of the Cbf1 HLH region is potentially positioned to interact with the highly basic N-terminal H3 tail, which is disordered and was not resolved by cryo-EM. Cbf1 interactions with histones appear to stabilize Cbf1 binding within the nucleosome resulting in the substantially slower dissociation rate from nucleosomes relative to DNA.

In addition to Cbf1-histone interactions, the Cbf1-nucleosome structure reveals that a PF can stabilize a partially unwrapped state of the nucleosome. This nucleosome “opening” likely provides access to additional DNA binding chromatin regulatory complexes. This is in line with previous structural studies that have revealed how a PF may increase local DNA accessibility. The Sox2-nucleosome and Sox11-nucleosome structures show that these PFs distort the nucleosomal DNA (Dodonova et al., 2020). In addition, the Oct4-Sox2-Nucleosome structures reveal that in combination these PFs disrupt histone-DNA contacts and stabilize partial DNA unwrapping from the histone octamer (Michael et al., 2020). However, in each of these cases, increased DNA accessibility results from induced DNA distortions upon binding. In contrast, Cbf1 interacts with both DNA and histones, which strengthen its stability relative to Cbf1 in complex with DNA. These additional interactions with the nucleosome enhance Cbf1’s ability to bind within the nucleosome and reduce its dissociation rate relative to DNA. This provides a molecular picture of the dissociation rate compensation mechanism (Donovan et al., 2019b).

Pioneer transcription factors possess two key properties (i) they access their target sites within nucleosomes efficiently, where they typically bind to nucleosomal DNA and bare DNA with similar affinities (Cirillo and Zaret, 1999; Cirillo et al., 1998; Donovan et al., 2019b; Fernandez Garcia et al., 2019; Sekiya et al., 2009; Soufi et al., 2014) and (ii) they initiate the formation of NDRs through nucleosome translocation and/or eviction (Bai et al., 2010; Yan et al., 2018). However, it is challenging to provide a quantitative and direct connection between these two PF properties. In this work, a direct connection is made between these two PF properties by investigating *in vivo* the Cbf1-Pho4 chimeras in budding yeast that we studied *in vitro*. Our results that Cbf1ΔN and Pho4ΔN CHLHZ, but not Cbf1ΔN PHLH and Pho4ΔN, efficiently repositions nucleosomes away from embedded Cbf1 motif indicate that the Cbf1 HLH region promotes nucleosome displacement activity *in vivo*. This is highly consistent with the *in vitro* measurement that the Cbf1 HLH region enables slow dissociation from the nucleosome and makes direct interactions with the histone octamer. However, our studies also show that Pho4ΔN can displace nucleosomes over a Pho4 motif. This is likely due to (i) Pho4ΔN has a very high affinity towards its consensus motif, and (ii) Pho4ΔN was expressed to a high level in this experiment. Indeed, even a prokaryotic TF can induce nucleosome displacement at high concentrations (Yan et al., 2018). It is possible that Cbf1ΔN and Pho4ΔN use two different mechanisms to invade nucleosomes: the former may employ an “active” mechanism where Cbf1ΔN directly docks onto nucleosomes and recruit downstream factors to reposition nucleosomes. The latter may use a more “passive” mechanism where Pho4ΔN takes advantage of intrinsic nucleosome dynamics to bind transiently exposed DNA and prevent nucleosome formation. Consistent with the “docking” state in the active mechanism, Pho4ΔN CHLHZ at lower concentration seems to bind within nucleosome − 4 and unwrap DNA but does not displace the nucleosome until it is highly expressed. In contrast, Pho4ΔN binding seems to be more correlated with nucleosome displacement. The detailed molecular pathways for the active vs passive nucleosome invasion require further investigation.

These combined results suggest a model (**Figure 6C**) where nucleosomes effectively block binding of Pho4 (a canonical TF) when expressed at low levels. However, Cbf1 (a PF), even at low expression levels, is able to bind a nucleosome-embedded motif through the dissociation rate compensation mechanism. Because of the extra interactions of Cbf1 with the histones, the nucleosome is not spontaneously repositioned. However, because Cbf1 has such a slow dissociation rate, there is time for it to recruit other chromatin modifying complexes most likely through the disordered N-terminal region, generating the large NDR that was observed in previous studies (Yan et al., 2018) (**Figure S19**). At higher levels of expression, both Cbf1 and Pho4 fully occupy their sites which results in spontaneous repositioning of the nucleosome and a smaller NDR that does not require either chromatin modifying complexes nor the disordered N-terminal region. The combined single molecule, cryo-EM structural and *in vivo* studies presented here provide significant molecular insight into how a PF interacts with the nucleosome to remain stably bound through the dissociation rate compensation mechanism to both directly reposition nucleosomes and to recruit other factors to generate large nucleosome depleted regions to begin the process of transcription activation.

## ACKNOWLEDGMENTS

We thank the members of the Poirier, Bai and Tan Labs for helpful discussions. We thank Joseph Cho for preparing cryo-EM grids and Jean-Paul Armache for helping with single particle data processing. This work was supported by NIH grants R01 GM121858 (to LB and MGP), R01 GM131626 and R35 GM139564 (to MGP), R35 GM139654 (to LB), R35 GM127034 (to ST), and T32 GM086252 (to BTD), by Estonian Research Council grant PUTJD906 (to PE) and by NSF grant 1715321 (to MGP). A portion of this research was supported by NIH grant U24GM129547 and performed at the PNCC at OHSU and accessed through EMSL (grid.436923.9), a DOE Office of Science User Facility sponsored by the Office of Biological and Environmental Research. We thank PNCC staff Rose Marie Haynes and Theo Humphreys for their expert assistance in cryo-EM data collection.

## STAR METHODS

### KEY RESOURCES TABLE

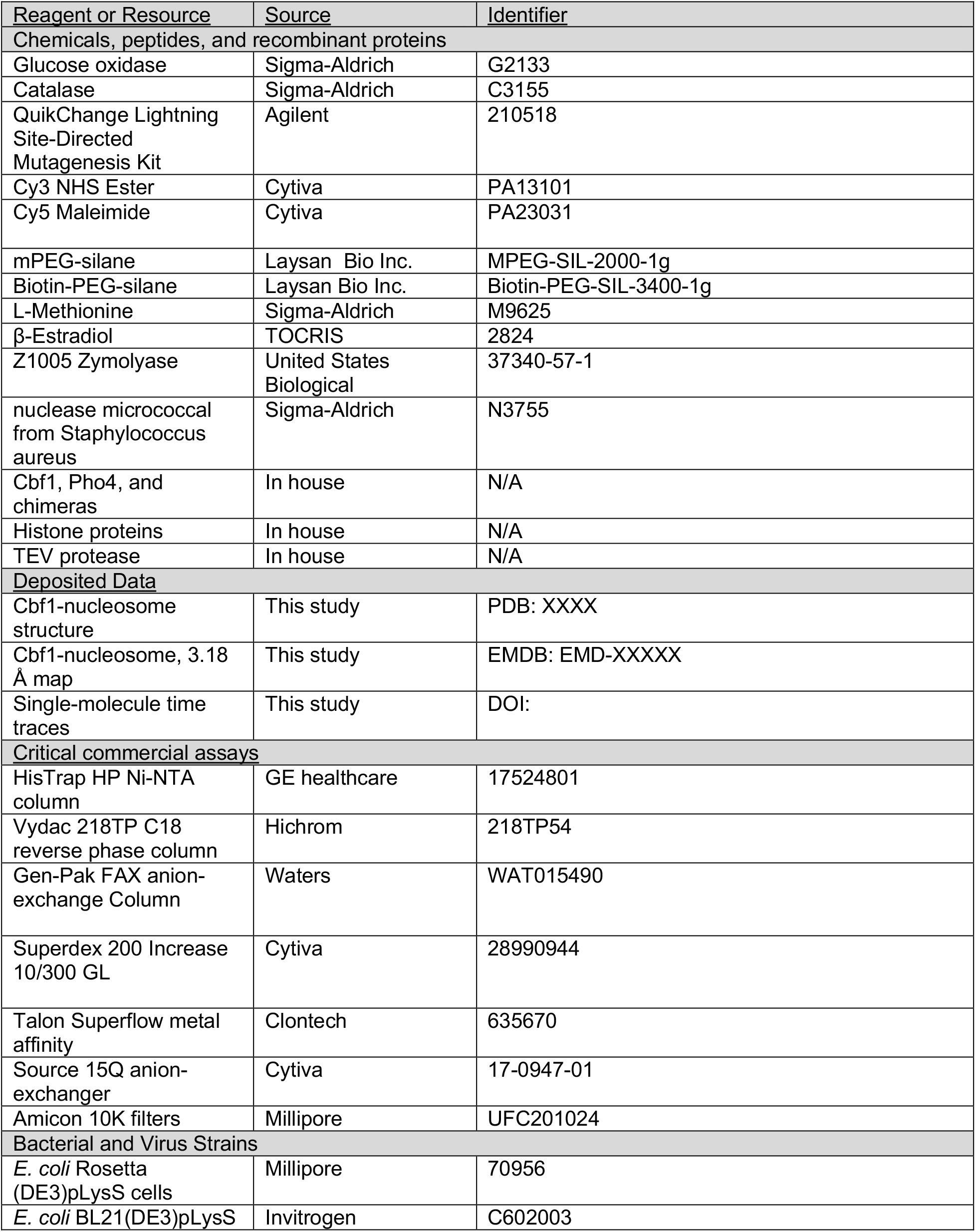
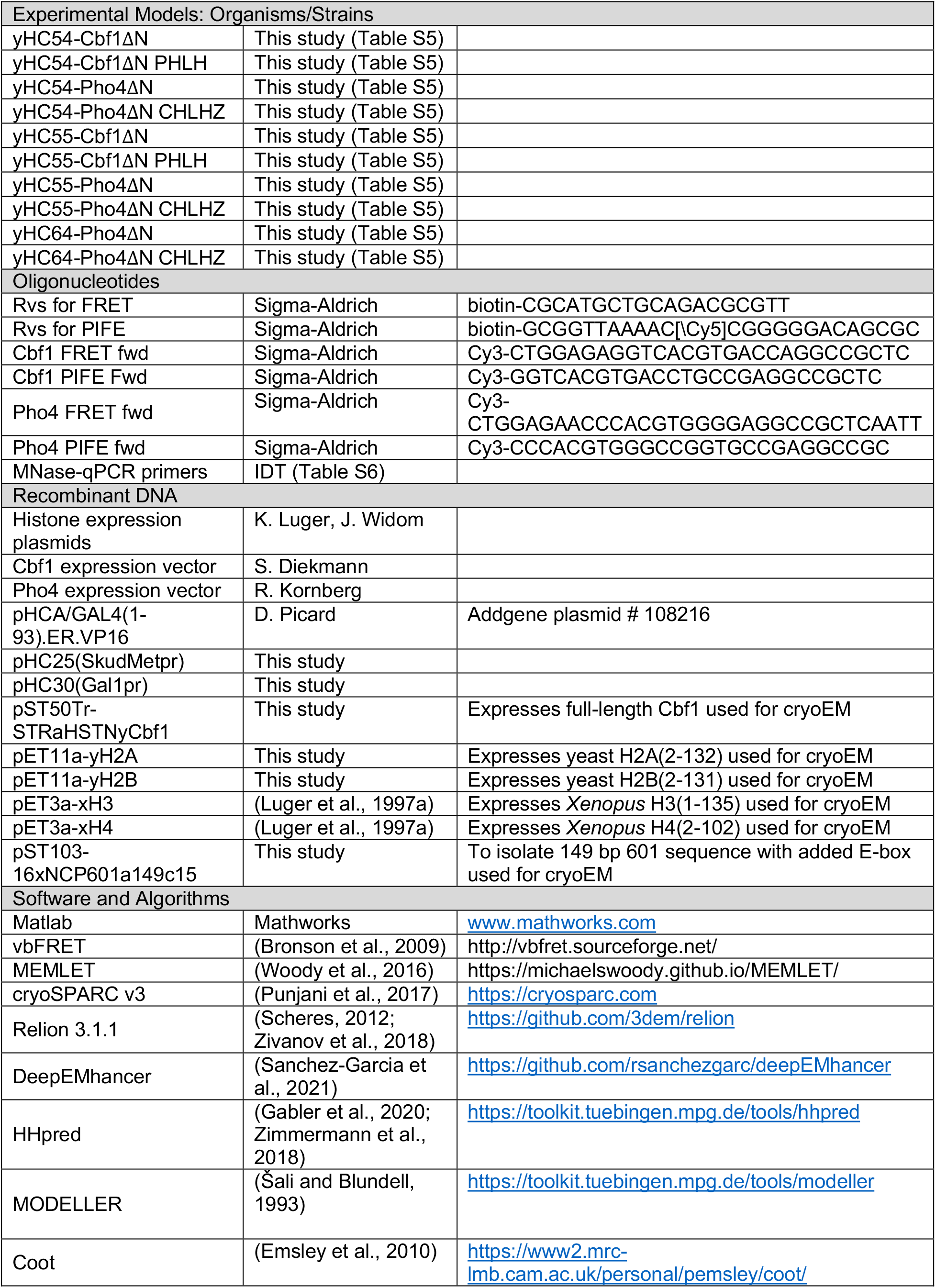
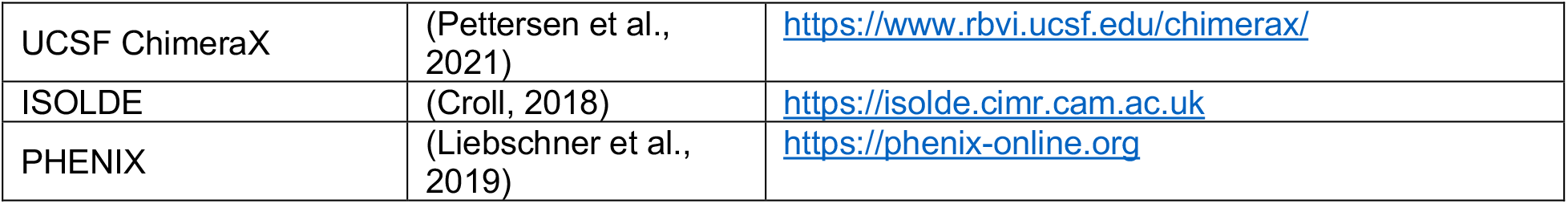

### RESOURCES AVAILABILITY

#### Lead Contact

Further information and request for resources and reagents should be directed and will be fulfilled by the Lead Contact, Michael G. Poirier

#### Materials Availability

Unique and stable reagents generated in this study are available upon request.

#### Data and Code Availability

A link to the time traces from single-molecule experiments will be available on Zenodo upon publication of the manuscript. These can be viewed using vbFRET in matlab (Bronson et al., 2009). The structural model and the cryo-EM map of Cbf1-nucleosome complex are available as of the date of publication at PDB XXXX and EMD-XXXXX, respectively.

### METHODS DETAILS

#### Preparation of Cbf1, Cbf1ΔN, Pho4ΔN CHLHZ

Cbf1 cloned into the pET28 expression vector was a gift from S Diekmann and expresses the Cbf1 coding sequence from the S288C strain of budding yeast. The expression vectors for Cbf1ΔN, and Pho4ΔN CHLHZ were prepared by site directed mutagenesis (Agilent) using the Cbf1 WT expression vector as a template. Proteins were expressed in BL21(DE3) cells (Invitrogen) and induced when the OD600 was between 0.4–0.6 with 0.5 mM IPTG for 4 hr at 37C. Cells were then pelted and frozen at −80 C until purification. Thawed cell pellets were resuspended in buffer A (50 mM Na2HP04 (pH 7.5), 300 mM NaCl, 5 mM imidazole, 10% glycerol, 1 mM PMSF, 20 ug/mL pepstatin, 20 ug/mL leupeptin) and lysed by sonication with cell debris removed by centrifugation (4 C, 23,000 x G, 20 min) and loaded directly onto a 5 mL HisTrap HP Ni-NTA column (GE Healthcare). Bound protein was washed with 40 mL buffer A, 120 mL buffer B (50 mM Na2HP04 (pH 7.5), 300 mM NaCl, 60 mM imidazole, 10% glycerol, 1 mM PMSF, 20 ug/mL pepstatin, 20 ug/mL leupeptin), and eluted with buffer C (50 mM Na2HP04 (pH 7.5), 300 mM NaCl, 340 mM imidazole, 10% glycerol, 1 mM PMSF, 20 ug/mL pepstatin, 20 ug/mL leupeptin). Pure fractions (as determined by SDS PAGE) were pooled, and imidazole was removed by dialysis into Buffer D (50 mM Na2HP04 (pH 7.5), 300 mM NaCl, 1 mM PMSF, 20 ug/mL pepstatin, 20 ug/mL leupeptin) in a 10 K amicon (Millipore). Concentration was determined by measuring absorbance at 280 nm. In all binding assays we report the concentration of protein dimer.

#### Preparation of Pho4, Pho4ΔN, Pho4 Zip, and Cbf1ΔN PHLH

The Pho4 expression vector was a gift from the R. Kornberg lab (Nagai et al., 2017). The Pho4ΔN, Pho4ΔN Zip, and Cbf1ΔN PHLH expression vectors were prepared by site directed mutagenesis (Agilent). All proteins were expressed in BL21(DE3) (invitrogen) by inducing at OD600 = 0.4-0.6 with 1 mM IPTG for 3 hours at 37 C. Cells were resuspended in 15 mL lysis buffer (20 mM Tris-HCL pH8, 500 mM NaCl, 10 mM imidazole, 0.2% Triton X-100, 5 mM DTT, 1 mM PMSF, 20μg/mL pepstatin, 20 μg/mL leupeptin, 2.1 mM benzamidine hydrochloride) per 600 mL culture and lysed by sonication. Cell debris was removed by centrifugation, loaded onto a 5 mL HisTrap HP Ni-NTA column (Cytiva), washed 150 mL wash buffer (20 mM Tris-HCL pH8, 500 mM NaCl, 5 mM imidazole, 0.02% Triton X-100, 5 mM DTT) followed by 150 mL wash of wash buffer without Triton X-100. Bound proteins were eluted in wash buffer with a gradient from 25-500 mM imidazole. Peak fractions were further purified with a superdex s200 10/300 (GE) size exclusion column equilibrated with storage buffer (40 mM HEPES-NaOH pH 7.4, 200 mM potassium acetate, 1 mM EDTA). Pure fractions, as determined by coomassie SDS PAGE, were pooled, concentrated, glycerol added to final concentration of 10% v/v, flash-frozen, and stored at −80C. In all binding assays we report the concentration of protein dimer.

#### Preparation of DNA molecules

DNA molecules were prepared by PCR using primers (Sigma, **Table S6**) from a plasmid containing the 601 nucleosome positioning sequence (NPS) (Lowary and Widom, 1998). Fluorescently labeled oligos were labeled at their 5’-ends with Cy3 or, in the case of the reverse primer for smPIFE, internally labeled at an amine modified dT with Cy5. Fluorescently labeled oligos were purified by HPLC with a 218TP C18 column (Grace/Vydac). After amplification, DNA molecules were purified using a MonoQ column (GE Healthcare).

#### Preparation of Histone Octamers

Human recombinant histones were purchased from histone source. Octamer reconstitution was performed with WT histones with the exception of H3(C110A) and H2A(K119C) which is required for labeling with Cy5 maleimide. The histone octamer was refolded by adding each of the histones together with H2A/H2B in 10% excess of H3/H4 as previously described (Luger et al., 1997b). The histone octamer was labeled with Cy5-maleimide (GE Healthcare) and purified with a superdex s200 10/300 (Cytiva) size exclusion column as previously described (Shimko et al., 2013).

#### Preparation of nucleosomes

Nucleosomes were reconstituted with Cy3-labeled DNA and purified Cy5-labeled histone octamer by double salt dialysis as previously described (Shimko et al., 2013). Dialyzed nucleosomes were loaded onto 5–30% sucrose gradients and purified by centrifugation on an Optima L-90 K Ultracentrifuge for 22 hrs (Beckman Coulter) with a SW-41 rotor. Sucrose fractions containing nucleosomes were collected, concentrated, and stored in 5x TE (pH 8) on ice.

#### Electrophoresis mobility shift assays

0.5 nM DNA or nucleosomes were incubated with TF in 10 mM Tris-HCl (pH 8), 130 mM NaCl, 10% glycerol, 0.0075% v/v Tween-20 for at least 5 min and then resolved by electrophoretic mobility shift assay (EMSA) with a 5% native polyacrylamide gel in 3x TBE. For very tight binders (i.e. Pho4 binding to DNA), the experiment was repeated with 0.2 nM DNA.

#### Ensemble PIFE measurements

Specific TF binding to Cy3-DNA was determined by protein-induced fluorescence enhancement (PIFE), in which Cy3 fluorescence increases upon protein binding. Fluorescence spectra were acquired with a Fluoromax4 fluorometer (Horiba) using an excitation wavelength of 510 nm. DNA was incubated for at least 5 min with varying concentrations of TF in 10 mM Tris-HCl (pH 8), 130 mM NaCl, 10% glycerol, and 0.0075% v/v Tween-20. DNA also contained a Cy5 fluorophore to normalize for small pipetting variabilities. Fluorescence spectra were analyzed using Matlab to determine the change in Cy3 fluorescence. For each assay, we selected a concentration of DNA that enabled clear visualization of binding. Binding affinity was determined using the concentration of unbound TF in the reaction.

#### Ensemble FRET measurements

TF binding to Cy3-Cy5 nucleosomes was measured as previously described (Li and Widom, 2004; Shimko et al., 2013). 0.2-0.5 nM nucleosomes were incubated for at least 5 min with varying concentrations of TF in 10 mM Tris-HCl (pH 8), 130 mM NaCl, 10% glycerol, and 0.0075% v/v Tween-20. Fluorescence emission spectra were acquired as previously described (Gibson et al., 2016). FRET efficiency was measured using the (Ratio)A method (Clegg, 1992).

#### Single molecule TIRF microscopy

The smTIRF microscope was built on an inverted IX73-inverted microscope (Olympus) as previously described (Roy et al., 2008). 532 nm and 638 nm diode lasers (Crystal Lasers) were used for Cy3 and Cy5 excitation. The excitation beams were expanded and then focused through a quartz prism (Melles Griot) at the surface of the quartz flow cell. A 1.3 N.A. silicone immersion objective (Olympus) was used to collect fluorescence, which was separately imaged onto an iXon3 EMCCD camera (Andor) with a custom-built emission path containing bandpass filters and dichroic beam splitters (Chroma Tech). Each video was acquired using Micro-Manager software (Open Imaging) (Tsuchida et al., 2014).

#### Flow Cell preparation

Flow cells were functionalized as previously described (Kinz-Thompson et al., 2013). Briefly, quartz microscope slides (Alfa Aesar) were sonicated in toluene and then ethanol, and then further cleaned by piranha (3:1 mixture of concentrated sulfuric acid to 50% hydrogen peroxide). Slides were washed in water and, once completely dry, incubated in 100 uM mPEG-Si and biotin-PEG-Si (Laysan Bio) overnight in anhydrous toluene. Functionalized quartz slides and coverslips were assembled into microscope flow cells where channels are defined with parafilm. Before each experiment, the flow cell was treated sequentially with 1 mg/ml BSA, 40 ug/ml streptavidin, and biotin-labeled DNA or nucleosomes.

#### Single-molecule fluorescence measurements of TF binding kinetics

Biotinylated sample molecules (DNA or nucleosomes) were allowed to incubate in the flow cell at room temperature for 5 min and then washed out with imaging buffer containing the desired TF concentration. The samples were first exposed to 638 nm excitation to determine the location of Cy5-labeled molecules and then to 532 nm excitation for both FRET and PIFE measurements. The imaging buffer for FRET experiments contained 10 mM Tris-HCl (pH 8), 130 mM NaCl, 10% glycerol, 0.5% v/v Tween-20, 0.1 mg/ml BSA, 2 mM Trolox, 0.0115% v/v COT, 0.012% v/v NBA, 450 ug/ml glucose oxidase (Sigma G2133) and 22 ug/ml catalase (Sigma C3155), while the imaging buffer for PIFE experiments contained 10 mM Tris-HCl (pH 8), 130 mM NaCl, 10% glycerol, 0.5% v/v Tween-20, 0.1 mg/ml BSA, 1% v/v BME, 450 ug/ml glucose oxidase and 22 ug/ml catalase.

Traces were selected manually. For smPIFE, we segregate molecules into three categories: (i) non-fluctuating molecules (typically about 30% of all traces), (ii) molecules that photobleach very quickly or have intensity above an empirically determined fluorescence value (intensity = 600) (approximately 20% of all traces), and (iii) molecules that are included for further analysis (about 40-50% of all traces). Category 3 molecules are then analyzed for interpretable fluctuations. The number of molecules analyzed for each experiment is included in supplemental information for each single-molecule experiment performed herein (typically about 20-50% of traces in category 3). For smFRET, we only analyzed traces that (a) FRET and (b) fluctuate. Again, we end up analyzing about 20-50% of all molecules that meet the initial criteria.

Single-molecule time series were fit to a two-state step function by the Hidden Markov Method using vbFRET (Bronson et al., 2009). From these idealized time series, we determined the dwell-time distributions of the TF bound and unbound states. Each distribution was analyzed using MEMLET to determine the best fit for the data and ultimately to obtain rate constants for the transitions between bound and unbound states (Woody et al., 2016). Log-likelihood ratio tests were performed to determine whether to fit dwell time distributions to either a single or double exponential distribution. Dwell time distributions were fit to the double-exponential model if 2 out of 3 replicates for all TF concentrations used in this experiment produced a P-value below 0.01.

#### Preparation of nucleosomes for single particle analysis

Nucleosomes containing 149 bp Widom 601 sequence with an added E-box motif (GGTCACGTGACC) positioned 9 bp downstream from the DNA end were prepared essentially as described previously (McGinty et al., 2016). We used the strategy of employing hybrid histone octamers containing yeast H2A and H2B with *Xenopus* H3 and H4 (Ranjan et al., 2013; Singh et al., 2019) to overcome problems producing large quantities of high-quality nucleosomes containing only yeast histones. This decision is justified by that the fact that yeast histones H3 and H4 are highly similar to *Xenopus* H3 and H4, and further by our structural observation that Cbf1 appears to contact only histone H2A and H2B proteins.

#### Expression and purification of Cbf1 for single particle analysis

The coding region of Cbf1(2-351) was cloned into pST50Tr vector (Tan et al., 2005), and the protein expressed in BL21(DE3)pLysS *Escherichia coli* cells by inducing with 0.5 mM isopropyl β-D-thiogalactopyranoside at 37°C. Cbf1 was purified from the crude cell extract by metal-affinity chromatography (Talon resin, Clontech) and digested with tobacco etch virus protease to remove the His_10_ affinity tag. The protein was further purified by SourceQ anion exchange chromatography (Cytiva). Pooled fractions were dialyzed against 10 mM HEPES pH 7.5, 200 mM NaCl, 10 mM β-mercaptoethanol, 0.1 mM PMSF, concentrated and supplemented with 20% v/v glycerol prior to flash freezing in liquid nitrogen and storage at −80°C.

#### Cbf1-nucleosome complex reconstitution for single particle analysis

Cbf1-nucleosome complexes were reconstituted by mixing Cbf1 with the E-box containing nucleosomes at a 2.5:1 molar ratio in 10 mM HEPES pH 7.5, 75 mM NaCl, 1 mM DTT. The complex was purified over a Superdex 200 Increase 10/300 size exclusion column (Cytiva) in the reconstitution buffer supplemented with 0.1 mM PMSF. Peak fractions were diluted and used for cryo-EM grid preparation.

#### Cryo-EM grid preparation and data collection

3 µl of Cbf1-nucleosome complex at 1 mg/ml was pipetted on a Quantifoil R 1.2/1.3 holey carbon grid (Electron Microscopy Sciences), blotted for 3 s with Vitrobot Mark IV (Thermo Scientific) at 4°C, 100% humidity, and plunge-frozen in liquid ethane at liquid nitrogen temperature. Grids were glow discharged for 45 s at 15 mA using an easiGlow device (PELCO) before sample application. The data was collected on a Krios G3i microscope (Thermo Scientific), operated at 300 keV, with a K3 direct electron detector camera (Gatan) using SerialEM (Mastronarde, 2005) at the Pacific Northwest Cryo-EM Center. 6339 movies were collected at a nominal magnification of 18,000x with super-resolution image pixel size of 0.644 Å/px (calibrated nominal pixel size 1.287 Å/px). The total dose per exposure was 50 e^-^/Å^2^, and each exposure was fractioned into 51 subframes. Data was recorded in the defocus range of −0.8 to −2.1 µm. The data collection parameters and model statistics are summarized in **Table S7**.

#### Single particle data processing

Initial data processing was performed in cryoSPARC v3 (Punjani et al., 2017). Super-resolution movies were motion-corrected with Patch Motion Correction and binned using 2x Fourier cropping. CTF values were estimated with Patch CTF Estimation. Initial model was created by picking particles with Blob Picker, the particles were 2D-classified, nucleosome-like classes were selected for *ab initio* reconstruction, and the best obtained model was used to generate templates for particle picking with Template Picker. Newly picked particles were again 2D-classified, noise-only classes were discarded, and *ab initio* reconstructions then generated from the obtained particles. Particles were further filtered through multiple rounds of Heterogeneous Refinement to remove junk and nucleosome-only particles, and to obtain classes of various Cbf1 conformations in relation to the nucleosome core. 1.9 million particles designated as Cbf1-nucleosome complexes were subjected to local (per-particle) motion correction (Punjani et al., 2017). To further discriminate between different Cbf1 conformations, particles classes from cryoSPARC’s Heterogeneous Refinement were transferred to Relion 3.1.1 (Scheres, 2012; Zivanov et al., 2018) for 3D classification without alignment (Scheres, 2016). Regularization parameter T was varied in the range of 100-400 and only the volume corresponding to Cbf1 was included in the mask. Several rounds of such classification were performed to obtain a set of conformationally homogenous particles. Final 3D refinement was performed with cryoSPARC’s Homogenous Refinement. The map was postprocessed using the deep learning algorithm DeepEMhancer (Sanchez-Garcia et al., 2021).

#### Model building and analysis

The initial model of Cbf1 was generated using HHpred (Gabler et al., 2020; Zimmermann et al., 2018) and MODELLER (Šali and Blundell, 1993). The best-fitting Cbf1 model and the crystal structure model of the nucleosome (PDB 3LZ0) were used for rigid-body fitting. The Cbf1-nucleosome model was built using Coot (Emsley et al., 2010) and UCSF ChimeraX (Pettersen et al., 2021) with the ISOLDE plugin (Croll, 2018). Real-space refinement was performed with the PHENIX suite (Liebschner et al., 2019) and validation with MolProbity (Williams et al., 2018). During refinement, secondary structure and Ramachandran restraints were employed. Density of Cbf1 residues 1-221 and 293-352 were not visible.

#### In vivo Nucleosome mapping

For methionine-induced strains, yeast cells were grown in synthetic complete media + 2% glucose+20xMethione (SCD+20xMet) overnight at 30°C until OD660 is 0.15, and then washed by ddH_2_O for three times. Washed cells were resuspended in SCD-Met medium and incubated for 2hr to fully induce the expression of target proteins. We imaged these cells and quantified the level of GFP-tagged proteins via MATLAB scripts (Zou and Bai, 2019). The subsequent nucleosome mapping follows previously described protocol (Bai et al., 2011). Briefly, cells were collected by centrifugation and then sequentially washed by cold water and 1M sorbitol. 1.5OD unit (OD660 x volume) of pre-washed cells were resuspended in 500ml of spheroplasting buffer (1M sorbitol and 0.5mM β-mercaptoethanol). 30µL of 0.5mg/ml of Zymolyase 100T was added to each sample, and incubated at room temperature for 6.5min. Cells were then centrifuged at 500g for 3min at 4°C and gently washed twice with 1M sorbitol. Each cell pellet was resuspended in 200uL of MNase digestion buffer (0.5mM spermidine, 10mM NaCl, 0.2M Tris-HCl pH7.5, 5mM MgCl2, 1mM CaCl2, 0.075% NP40, 1M sorbitol, and 1mM β-mercaptoethanol), and 5µL of 0.2U/µl MNase was added. Samples were incubated at 37°C for 8min, and then reactions were quenched by adding 20ul of quench buffer (5% SDS and 0.25M EDTA). Quenched samples were incubated at 65°C for 10min. Genomic DNA was extracted with 200µl of Phenol:Chloroform:Isoamyl Alcohol (25:24:1) and precipitated with ethanol. DNA pellets were resuspended in 400ul of TE and digested with 100µg RNase A. DNA was pellet down again with 1ml of ethanol and 6µl of 5M ammonium acetate. Mono-nucleosomal fragments were then gel purified with a Genejet kit and subjected to stacking qPCR (see **Table S8** for primers). Nucleosome occupancy was normalized to a well-position nucleosome in the terminator of *EXO84*.

For β-estradiol-induced strains, cells were grown in SCD+10xMet overnight at 30°C until OD660 is 0.15. β-estradiol was added to cell culture to desired concentrations and incubated for 1.5hr. We analyzed these cells with the same imaging and MNase-qPCR assay as described above.

## SUPPLEMENTAL FIGURES

**Figure S1.**
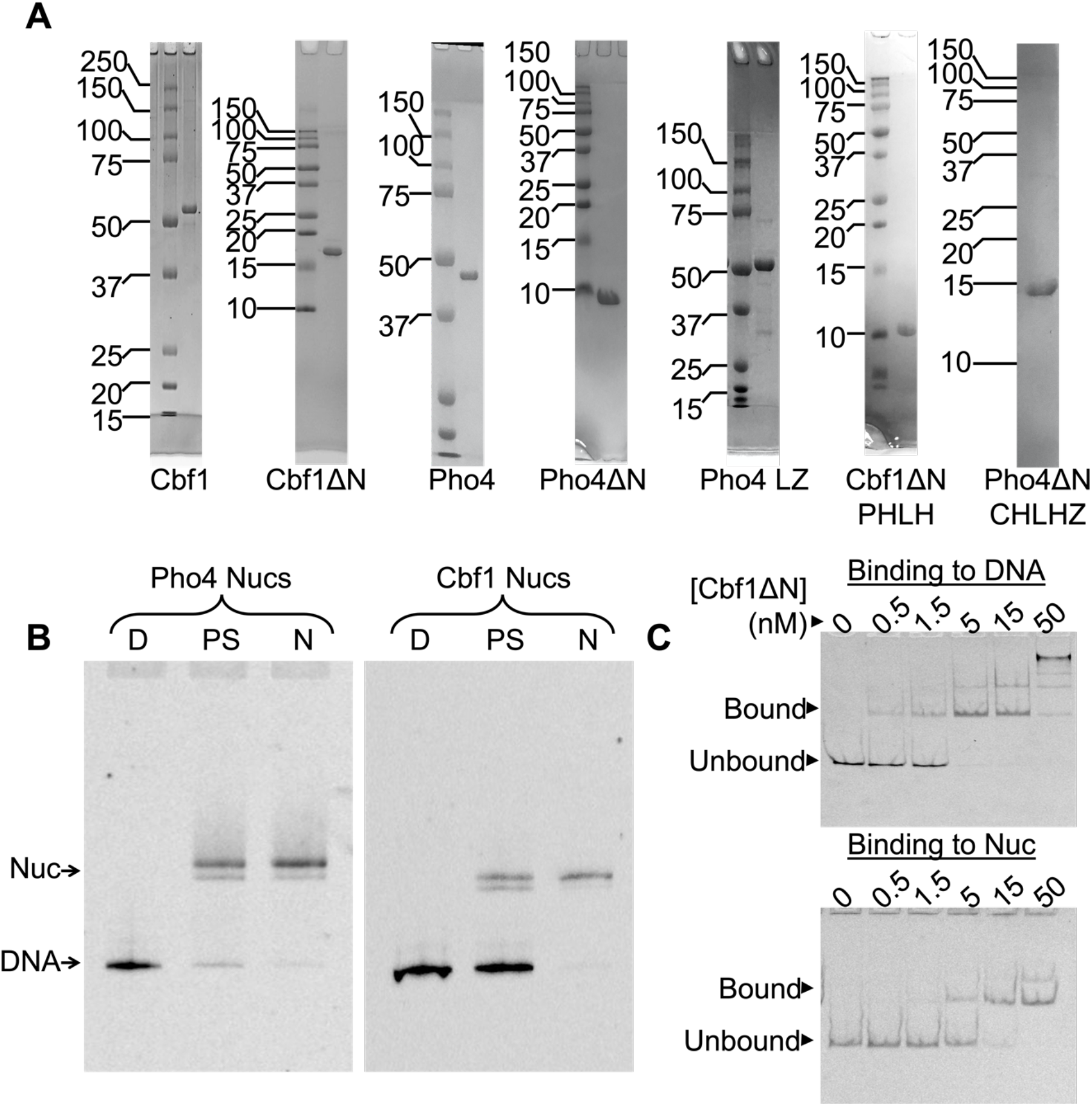
Protein and nucleosome samples prepared for this study. A) SDS gels of purified proteins used in this study. B) Representative native gels of nucleosomes used in this study, D = DNA, PS = pre-sucrose gradient purification, N = final nucleosomes. C) Electrophoresis mobility shift assays (EMSAs) to detect Cbf1ΔN binding to DNA (top) and nucleosomes (bottom).

**Figure S2.**
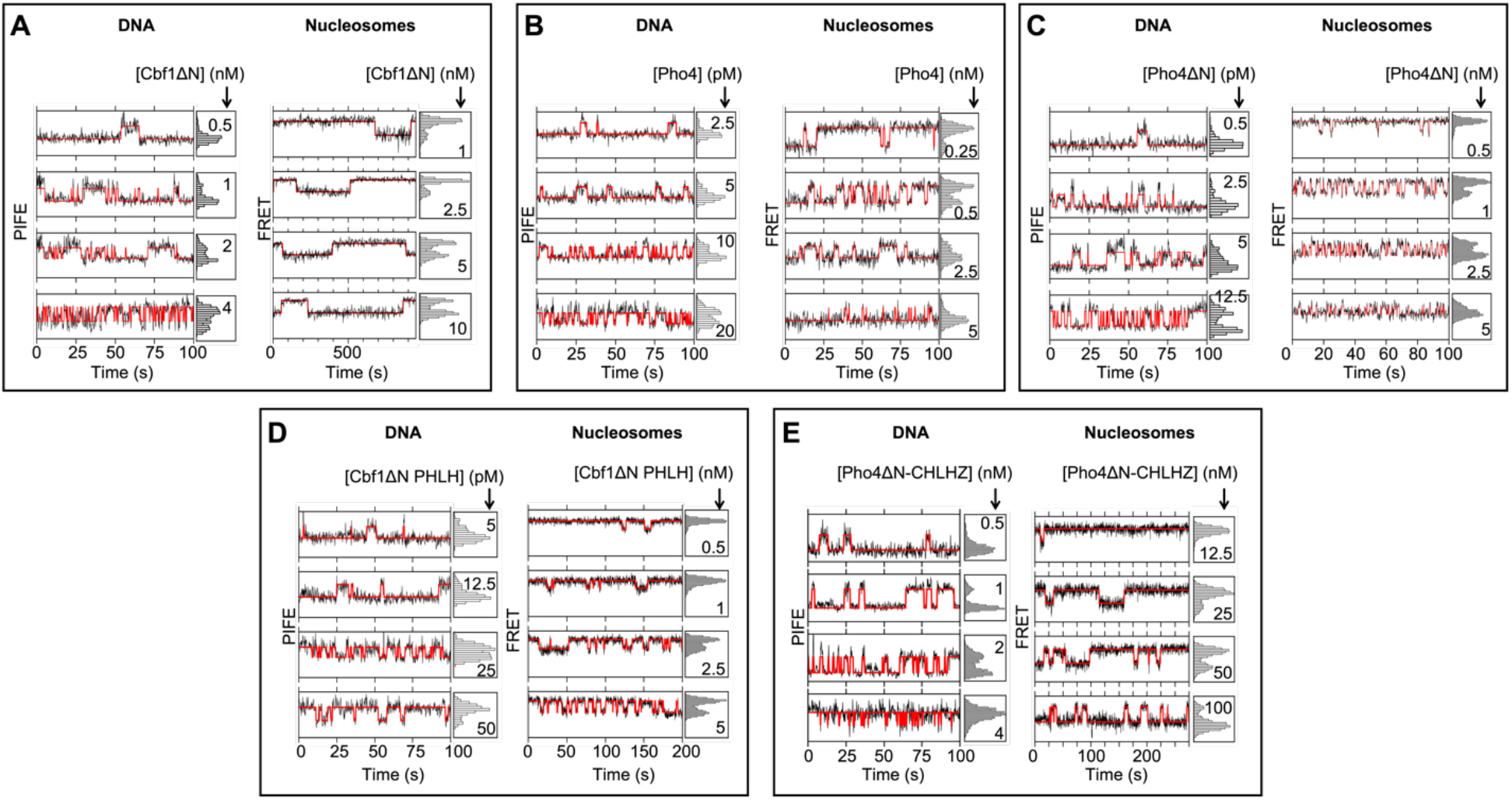
Representative example traces from smPIFE and smFRET measurements of (**A**) Cbf1ΔN, (**B**) Pho4, (**C**) Pho4ΔN, (**D**) Cbf1ΔN PHLH, and (**E**) Pho4ΔN CHLHZ binding to DNA and nucleosomes, respectively.

**Figure S3.**
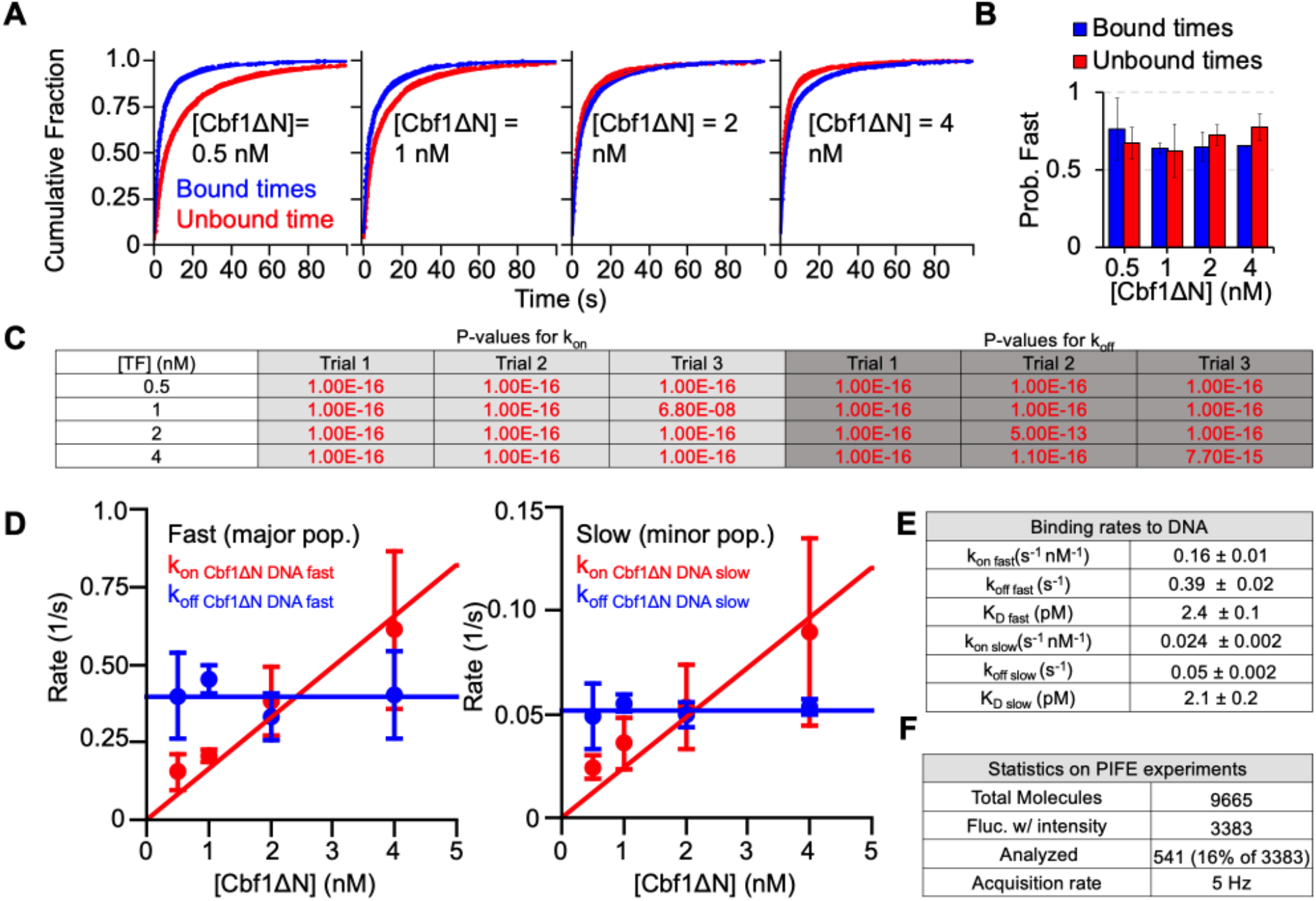
Single molecule characterization of Cbf1ΔN binding to DNA. (**A**) Cumulative sum distributions of bound (blue) and unbound dwell times (red) at all concentrations assayed. (**B**) The probability of bound (blue bars) or unbound (red bars) dwell times belonging to the fast population for all concentrations assayed. (**C**) Log-likelihood ratio tests were performed to determine whether to fit dwell time distributions to either a single or double exponential distribution. For each test, the null hypothesis was that the distribution follows a single-exponential decay. The null-hypothesis was rejected for P-values below 0.01. Rejected tests are indicated in red. Dwell time distributions were fit to the double-exponential model if 2 out of 3 replicates for all TF concentrations used in this experiment produced a P-value below 0.01. In this experiment, both the bound time and unbound time cumulative sums fit best to a double exponential. The resulting fits are included in (A). (**D**) Major (left) and minor (right) binding and dissociation rates for increasing concentrations of Cbf1ΔN. (**E**) Overall binding rate constants and dissociation rates determined from the fits in (D). (**F**) Statistics on the total number of molecules assayed. “Fluctuate with intensity” refers to the number of molecules with intensity values below 600 which was the empirically determined optimal intensity range for single-molecule PIFE.

**Figure S4.**
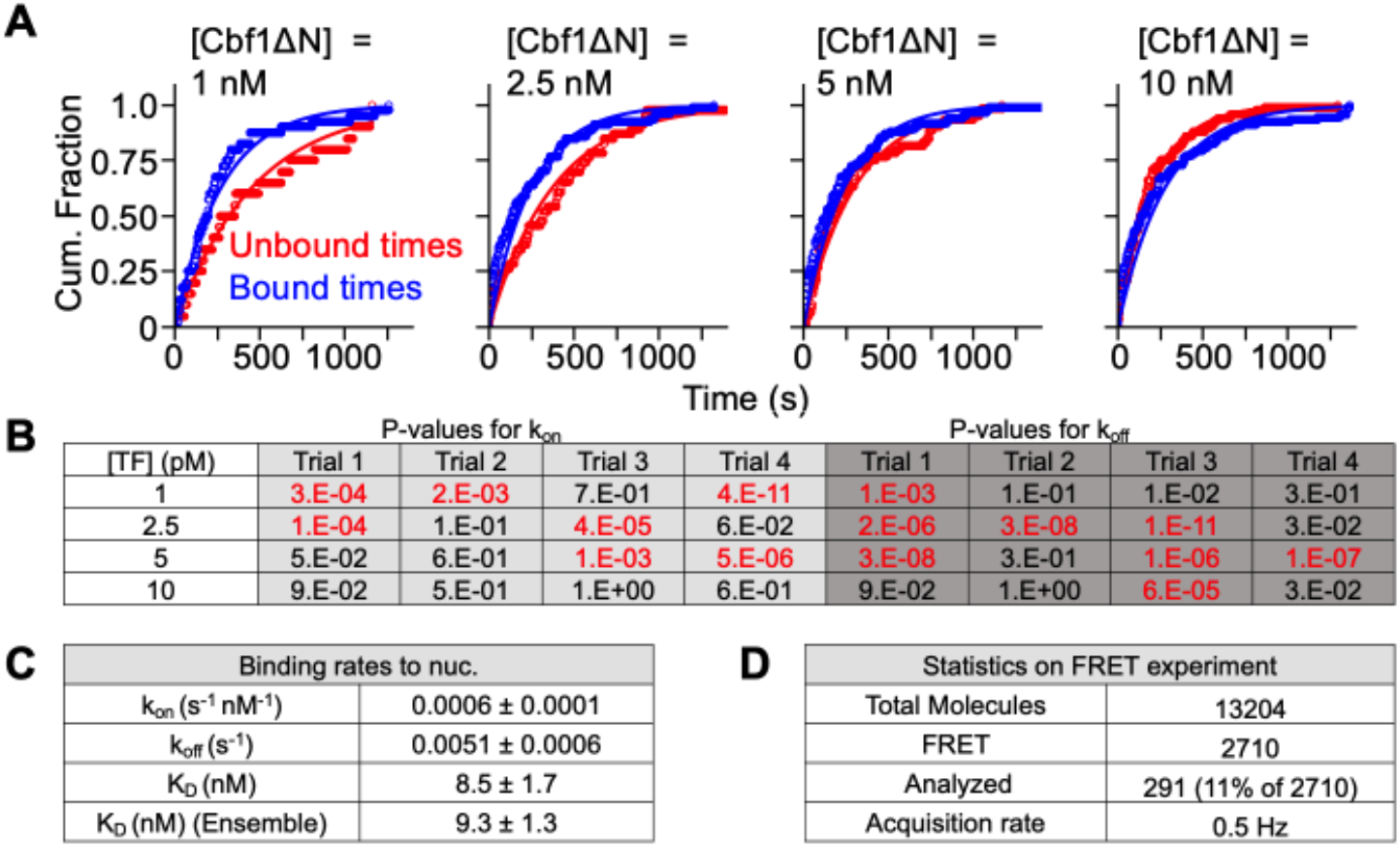
Single molecule characterization of Cbf1ΔN binding to Nucleosomes. (**A**) Cumulative sum distributions of bound (blue) and unbound dwell times (red) at all concentrations assayed. (**B**) Log-likelihood ratio tests were performed to determine whether to fit dwell time distributions to either a single or double exponential distribution. For each test, the null hypothesis was that the distribution follows a single-exponential decay. The null-hypothesis was rejected for P-values below 0.01. Rejected tests are indicated in red. Dwell time distributions were fit to the double-exponential model if 2 out of 3 replicates for all TF concentrations used in this experiment produced a P-value below 0.01. In this experiment, both the bound time and unbound time cumulative sums fit best to a single exponential. The resulting fits are included in (A). (**C**) The binding rate constant and dissociation rate determined from increasing concentrations of TF. (**D**) Statistics on the number of total molecules assayed. “FRET” refers to the number of molecules that emit from both Cy3 and a Cy5 fluorophores upon excitation at 532 nm and undergo fluctuations in FRET efficiency.

**Figure S5:**
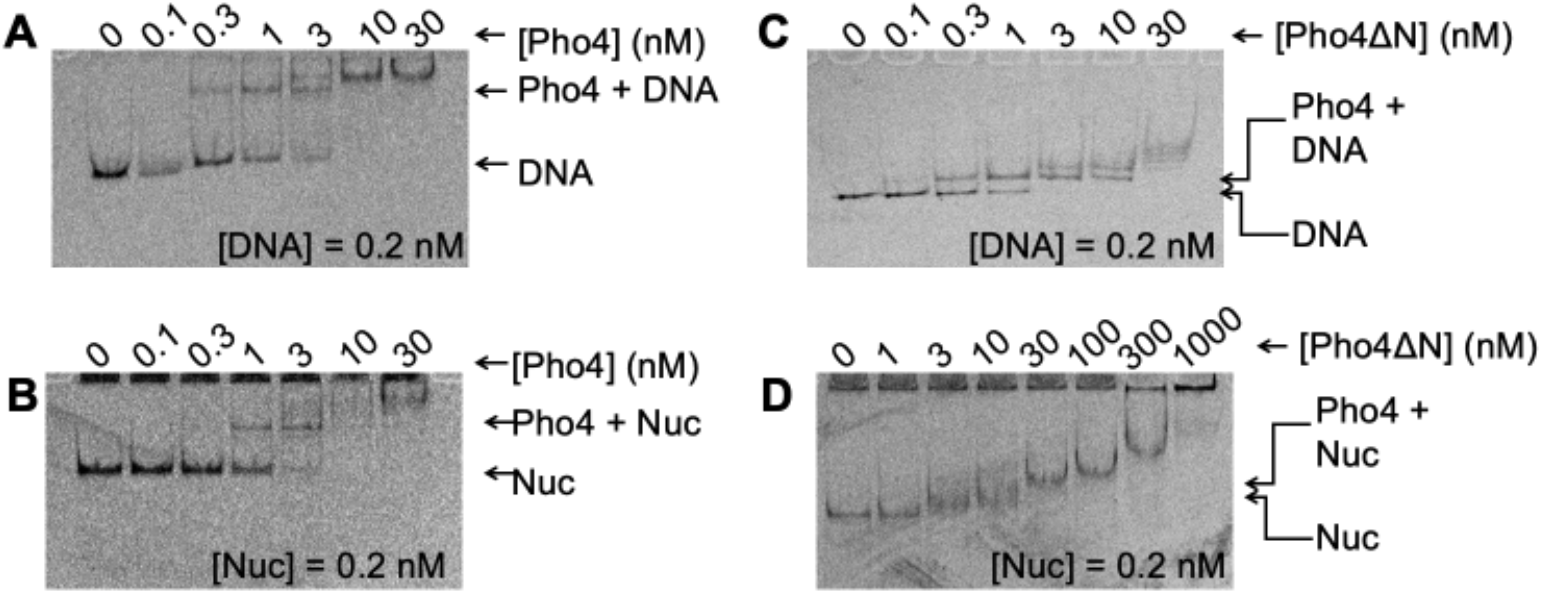
Characterization of Pho4 and Pho4ΔN binding to DNA and nucleosomes by EMSA. (**A**) Electrophoresis mobility shift assay (EMSA) of Pho4 binding to DNA. (**B**) EMSA of Pho4 binding to nucleosomes. (**C**) EMSA of Pho4ΔN binding to DNA. (**D**) EMSA of Pho4ΔN binding to nucleosomes.

**Figure S6.**
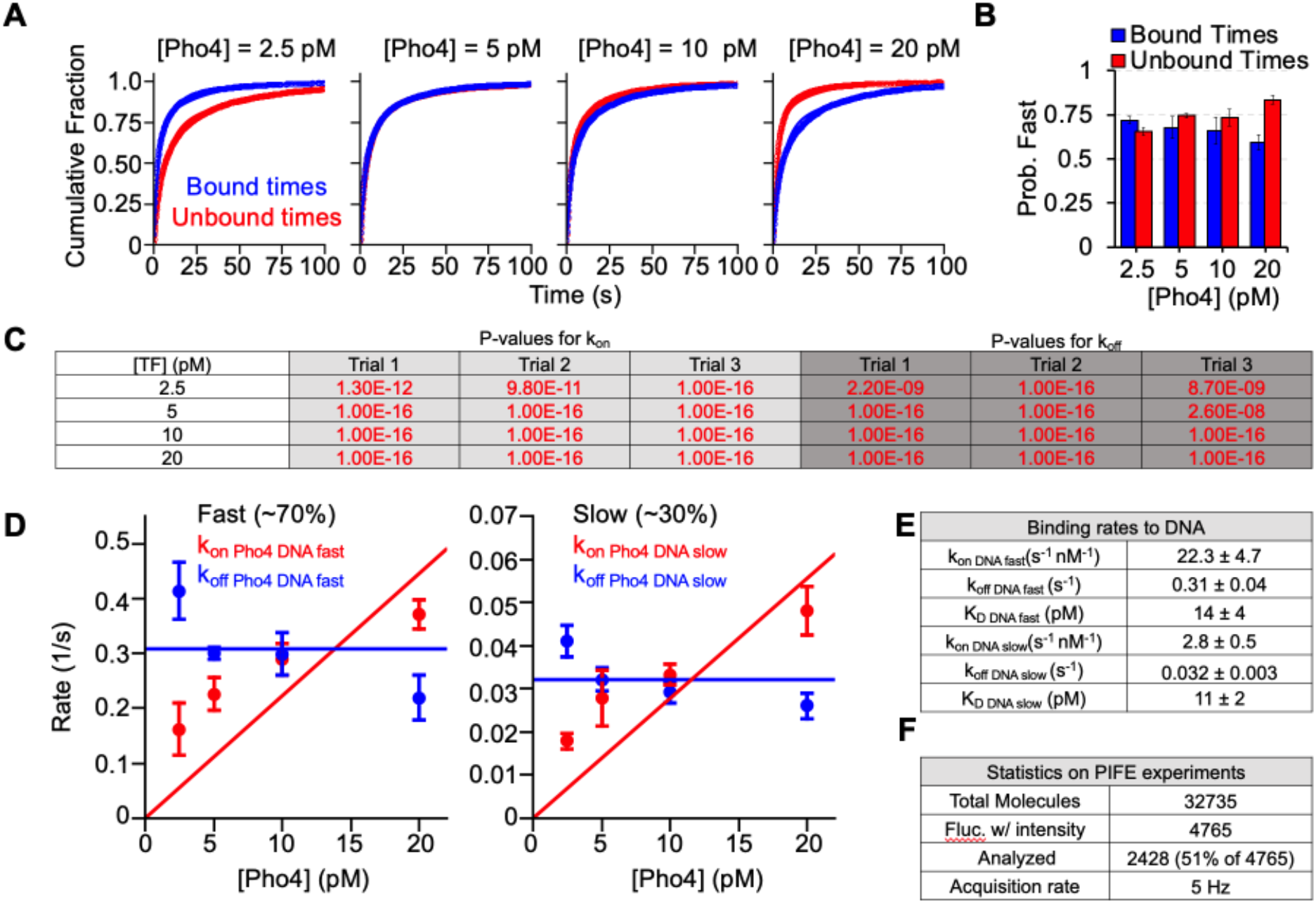
Single molecule characterization of Pho4 binding to DNA. (**A**) Cumulative sum distributions of bound (blue) and unbound dwell times (red) at all concentrations assayed. (**B**) The probability of bound (blue bars) or unbound (red bars) dwell times belonging to the fast population for all concentrations assayed. (**C**) Log-likelihood ratio tests were performed to determine whether to fit dwell time distributions to either a single or double exponential distribution. For each test, the null hypothesis was that the distribution follows a single-exponential decay. The null-hypothesis was rejected for P-values below 0.01. Rejected tests are indicated in red. Dwell time distributions were fit to the double-exponential model if 2 out of 3 replicates for all TF concentrations used in this experiment produced a P-value below 0.01. In this experiment, both the bound time and unbound time cumulative sums fit best to a double exponential. The resulting fits are

**Figure S7.**
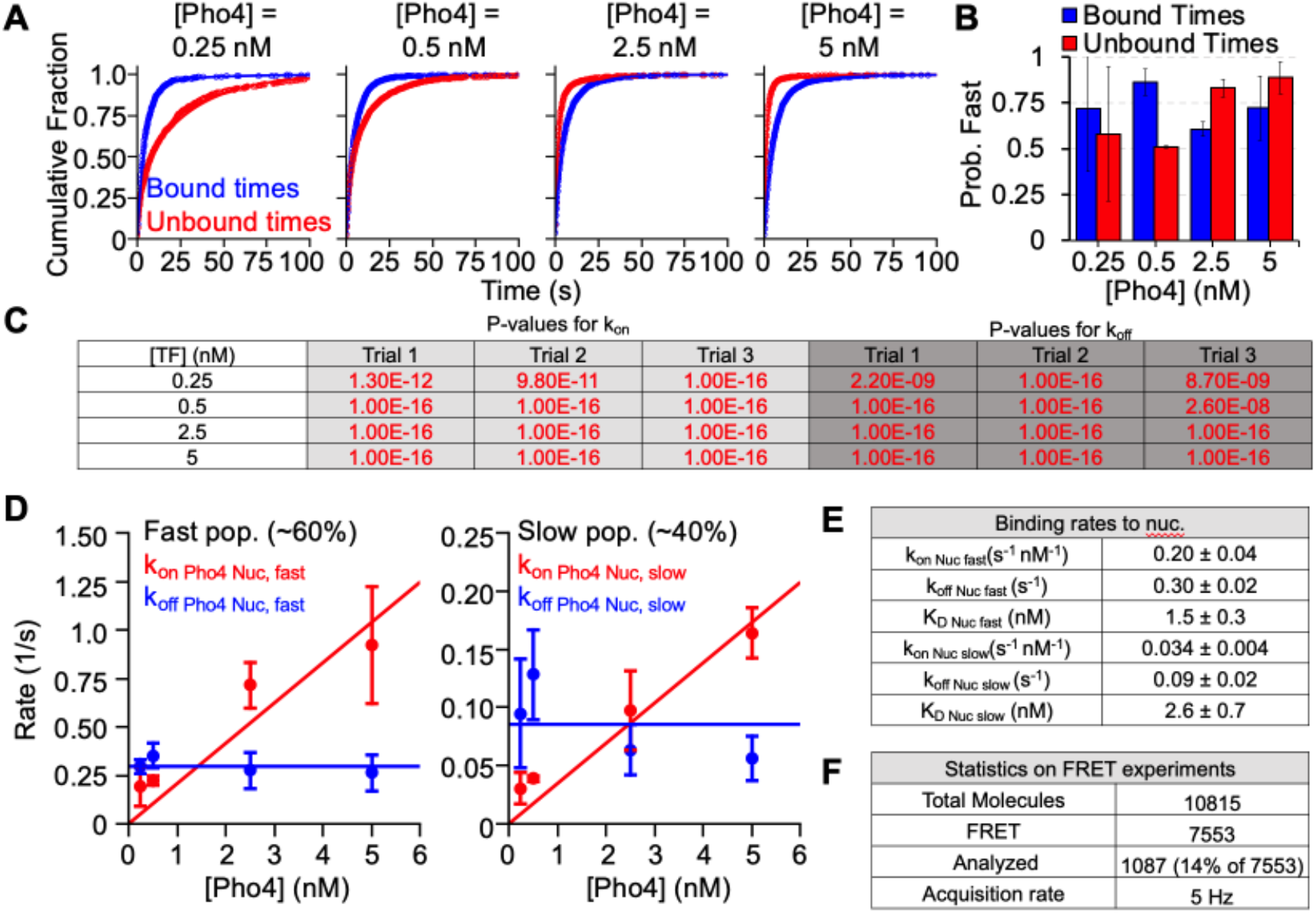
Single molecule characterization of Pho4 binding to nucleosomes. (**A**) Cumulative sum distributions of bound (blue) and unbound dwell times (red) at all concentrations assayed. (**B**) The probability of bound (blue bars) or unbound (red bars) dwell times belonging to the fast population for all concentrations assayed. (**C**) Log-likelihood ratio tests were performed to determine whether to fit dwell time distributions to either a single or double exponential distribution. For each test, the null hypothesis was that the distribution follows a single-exponential decay. The null-hypothesis was rejected for P-values below 0.01. Rejected tests are indicated in red. Dwell time distributions were fit to the double-exponential model if 2 out of 3 replicates for all TF concentrations used in this experiment produced a P-value below 0.01. In this experiment, both the bound time and unbound time cumulative sums fit best to a double exponential. The resulting fits are included in (A). (**D**) Major (left) and minor (right) binding and dissociation rates for increasing concentrations of TF. (**E**) Binding and dissociation rates determined from the fits in (D). (**F**) Statistics on the number of total molecules assayed. “FRET” refers to the number of molecules that emit from both Cy3 and a Cy5 fluorophores upon excitation at 532 nm and undergo fluctuations in FRET efficiency.

**Figure S8.**
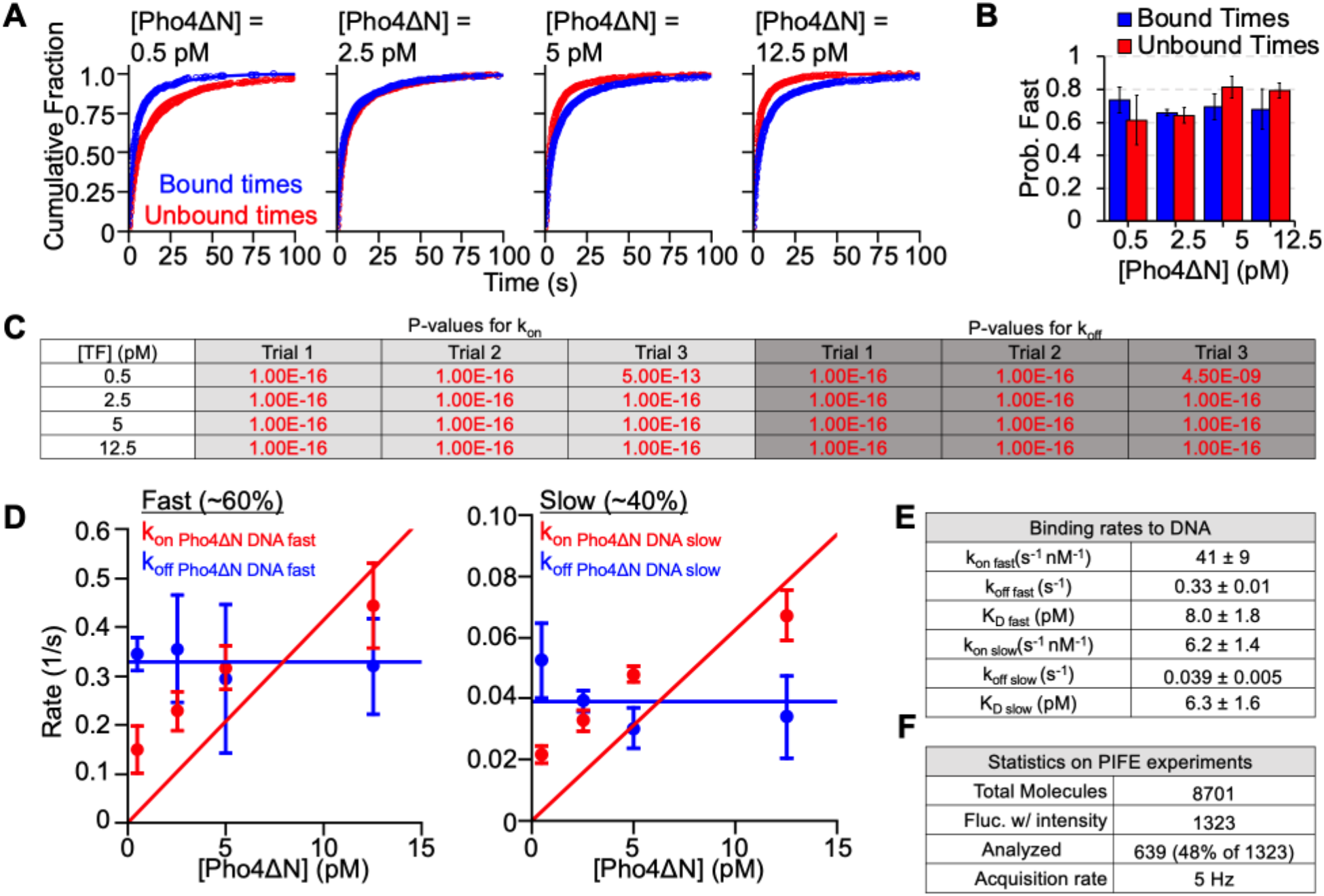
Single molecule characterization of Pho4ΔN binding to DNA. (**A**) Cumulative sum distributions of bound (blue) and unbound dwell times (red) at all concentrations assayed. (**B**) The probability of bound (blue bars) or unbound (red bars) dwell times belonging to the fast population for all concentrations assayed. (**C**) Log-likelihood ratio tests were performed to determine whether to fit dwell time distributions to either a single or double exponential distribution. For each test, the null hypothesis was that the distribution follows a single-exponential decay. The null-hypothesis was rejected for P-values below 0.01. Rejected tests are indicated in red. Dwell time distributions were fit to the double-exponential model if 2 out of 3 replicates for all TF concentrations used in this experiment produced a P-value below 0.01. In this experiment, both the bound time and unbound time cumulative sums fit best to a double exponential. The resulting fits are

**Figure S9.**
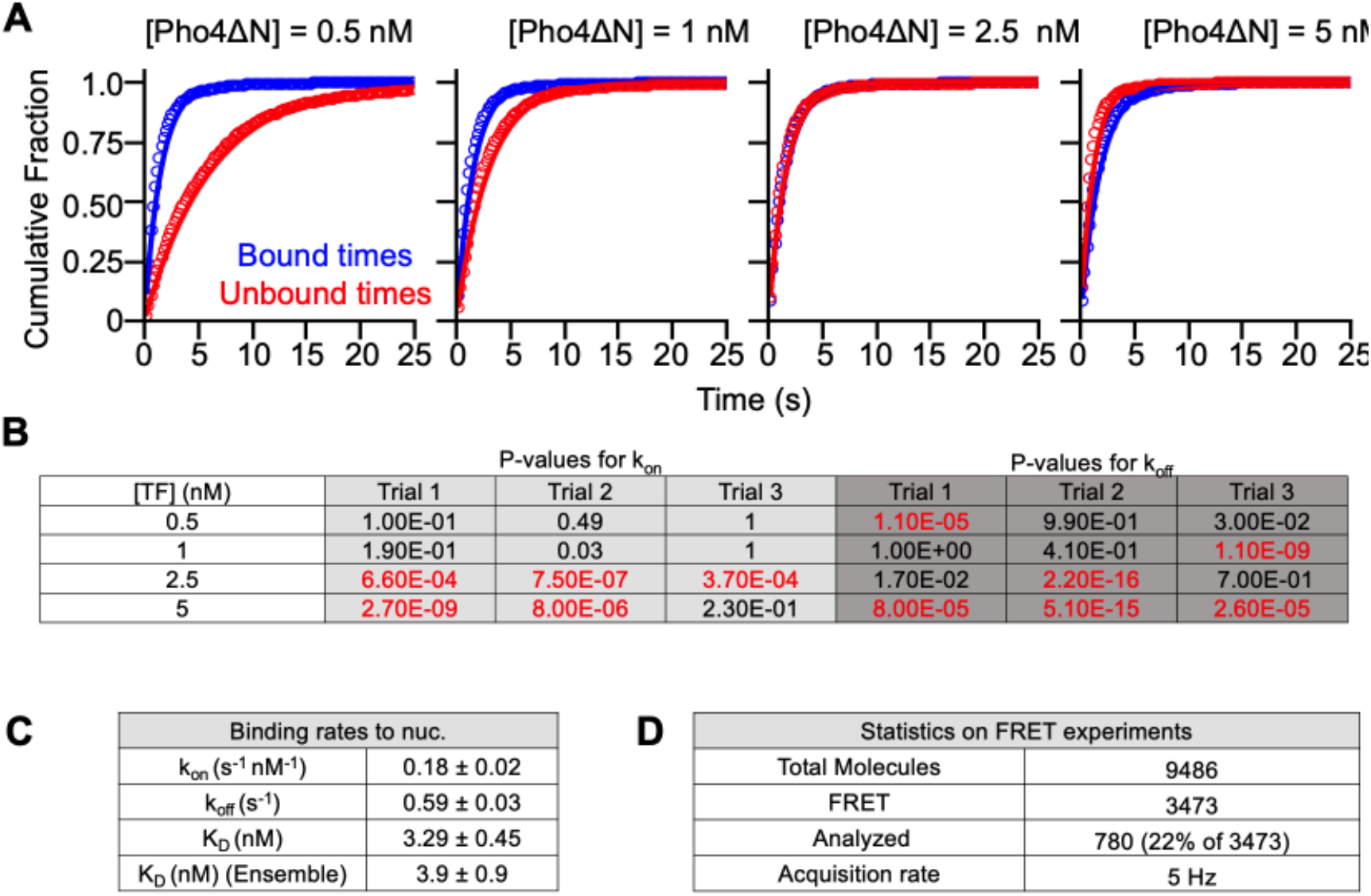
Single molecule characterization of Pho4ΔN binding to Nucleosomes. (**A**) Cumulative sum distributions of bound (blue) and unbound dwell times (red) at all concentrations assayed. (**B**) Log-likelihood ratio tests were performed to determine whether to fit dwell time distributions to either a single or double exponential distribution. For each test, the null hypothesis was that the distribution follows a single-exponential decay. The null-hypothesis was rejected for P-values below 0.01. Rejected tests are indicated in red. Dwell time distributions were fit to the double-exponential model if 2 out of 3 replicates for all TF concentrations used in this experiment produced a P-value below 0.01. In this experiment, both the bound time and unbound time cumulative sums fit best to a single exponential. The resulting fits are included in (A). (**C**) The binding rate constant and dissociation rate determined from increasing concentrations of TF. (**D**) Statistics on the number of total molecules assayed. “FRET” refers to the number of molecules that emit from both Cy3 and a Cy5 fluorophores upon excitation at 532 nm and undergo fluctuations in FRET efficiency.

**Figure S10.**
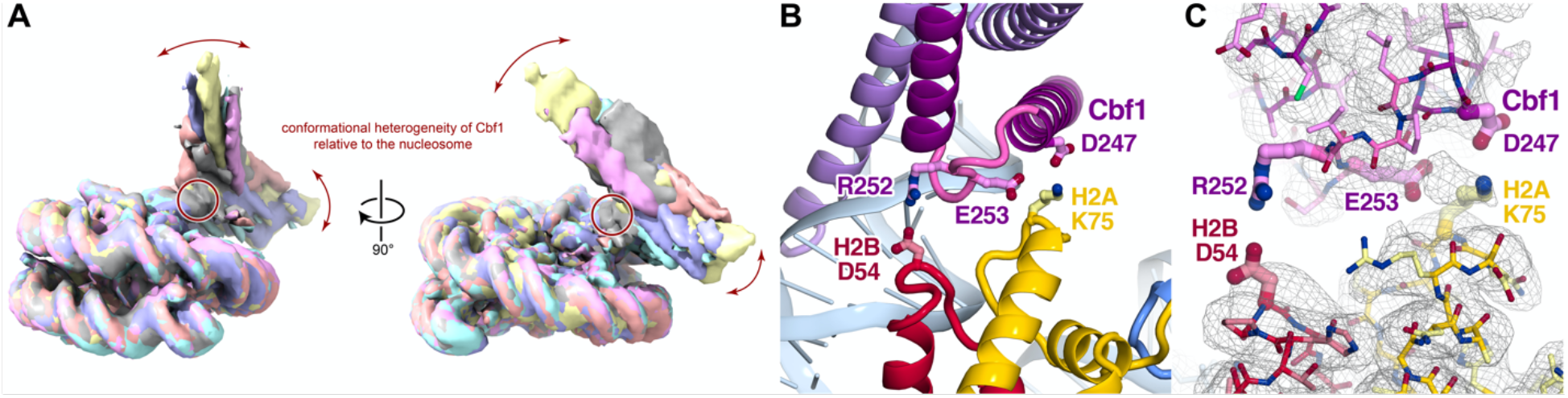
Cryo-EM analysis of Cbf1-nucleosome complex. **(A)** Overlayed reconstructions of 3D classes illustrate the continuous movement of Cbf1 and the unwrapped DNA in multiple directions (red arrows). This movement pivots around the cluster of electrostatic interactions between Cbf1 and the histone core (red circle). The reconstructions were obtained by heterogeneous refinement of 1.7M particles into 6 classes in cryoSPARC, each color representing a distinct 3D class. **(B)** Electrostatic interactions in the Cbf1-histone interface: Cbf1(D247) on helix1 and Cbf1(E253) in the HLH loop both interact with H2A(K75), while Cbf1(R252) interacts with H2B(D54). **(C)** Similar view as S10C overlaid with map density.

**Figure S11:**
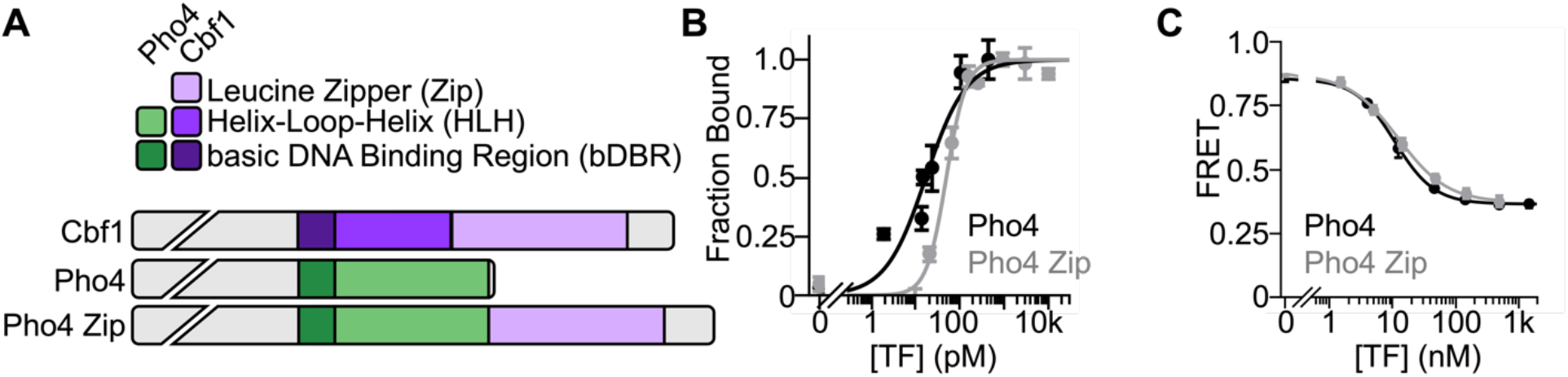
Addition of the leucine zipper onto Pho4 does not impact DNA or nucleosome binding. (**A**) Domain diagrams of Cbf1, Pho4, and the Pho4 Zip chimera, in which the Cbf1 leucine zipper is inserted onto the C-terminus of Pho4. (**B**) Ensemble PIFE measurements of Pho4 and Pho4 Zip binding at increasing concentrations to DNA. The x-axis displays the estimated concentration of unbound TF. In both experiments, we measure stoichiometric binding to DNA. (**C**) Ensemble FRET efficiency measurements of Pho4 and Pho4 Zip binding at increasing concentrations to nucleosomes. We measure no difference between Pho4 and Pho4 Zip binding affinity to nucleosomes when titrating either TF and measuring the corresponding FRET efficiency, S_1/2 Pho4 Nuc_ = 1.1 ± 0.1 nM, S_1/2 Pho4 Zip Nuc_ = 1.2 ± 0.1 nM.

**Figure S12:**
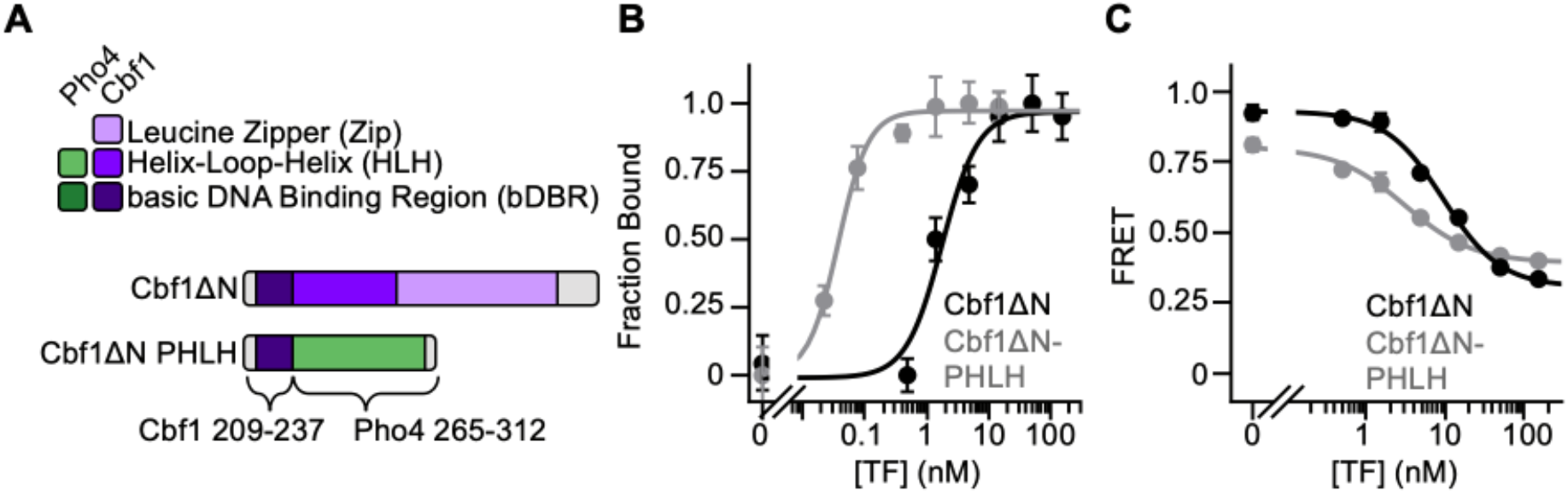
Ensemble fluorescence characterization of Cbf1ΔN PHLH. (**A**) Domain diagrams of Cbf1ΔN and Cbf1ΔN PHLH in which the Pho4 dimerization domain (Pho4 residues 265-312) is inserted after Cbf1 residues 209-237. (**B**) Ensemble PIFE measurements of Cbf1ΔN and Cbf1ΔN PHLH binding at increasing concentrations to DNA (S_1/2 Cbf1ΔN DNA_ = 1.8 ± 0.6 nM, S_1/2 Cbf1ΔN PHLH DNA_ = 37 ± 5 pM). The x-axis represents the estimated concentration of unbound protein. (**C**) FRET efficiency measurements of Cbf1ΔN and Cbf1ΔN PHLH binding at increasing concentrations to nucleosomes (S_1/2 Cbf1ΔN Nuc_ = 9.3 ± 1.3 nM, S_1/2 Cbf1ΔN PHLH Nuc_ = 3 ± 0.2 nM).

**Figure S13.**
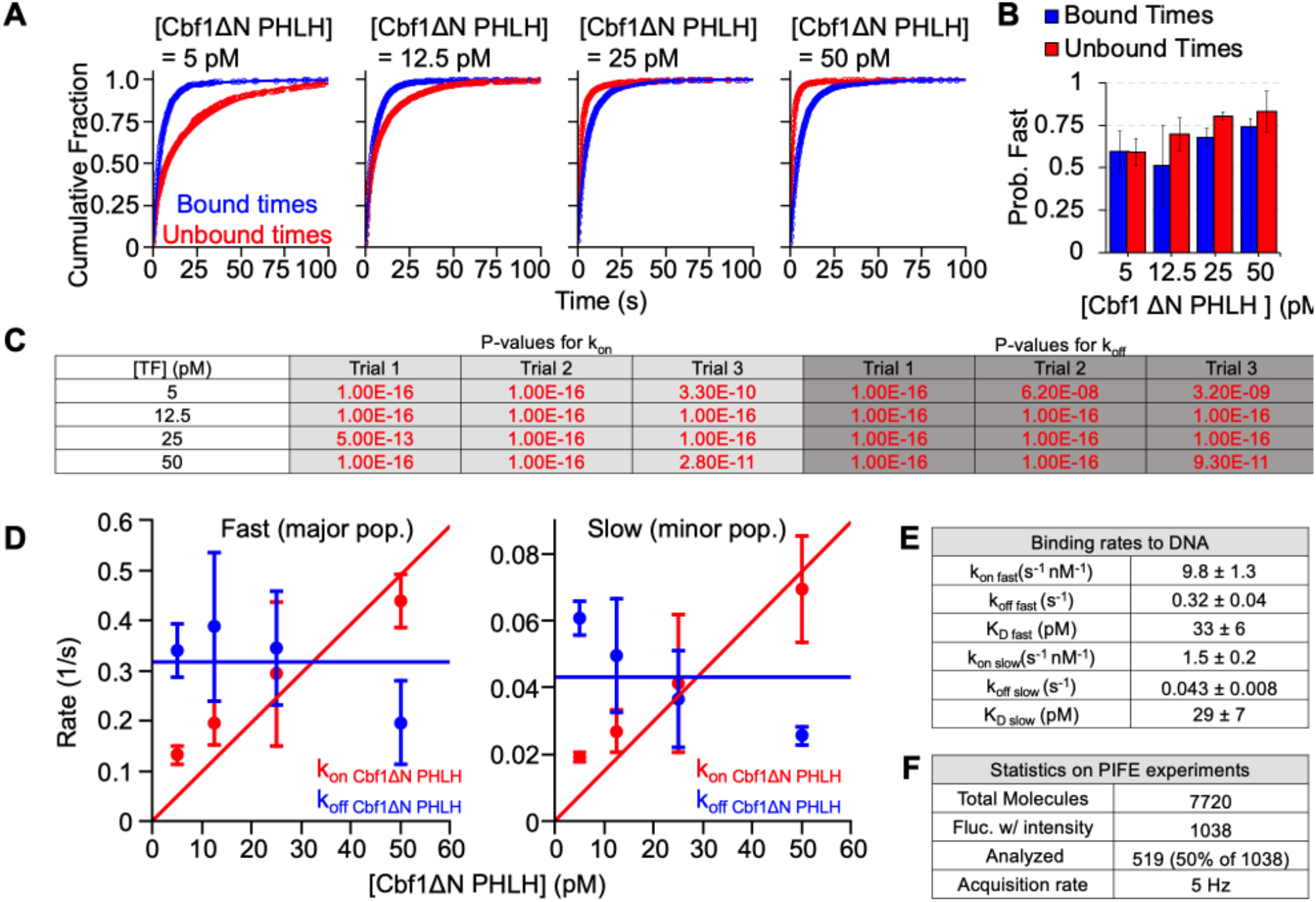
Single molecule characterization of Cbf1ΔN PHLH binding to DNA. (**A**) Cumulative sum distributions of bound (blue) and unbound dwell times (red) at all concentrations assayed. (**B**) The probability of bound (blue bars) or unbound (red bars) dwell times belonging to the fast population for all concentrations assayed. (**C**) Log-likelihood ratio tests were performed to determine whether to fit dwell time distributions to either a single or double exponential distribution. For each test, the null hypothesis was that the distribution follows a single-exponential decay. The null-hypothesis was rejected for P-values below 0.01. Rejected tests are indicated in red. Dwell time distributions were fit to the double-exponential model if 2 out of 3 replicates for all TF concentrations used in this experiment produced a P-value below 0.01. In this experiment, both the bound time and unbound time cumulative sums fit best to a double exponential. The resulting fits are included in (A). (**D**) Major (left) and minor (right) binding and dissociation rates for increasing concentrations of TF. (**E**) Overall binding rate constants and dissociation rates determined from the fits in (D). (**F**) Statistics on the total number of molecules assayed. “Fluctuate with intensity” refers to the number of molecules with intensity values below 600 which was the empirically determined optimal intensity range for single-molecule PIFE.

**Figure S14.**
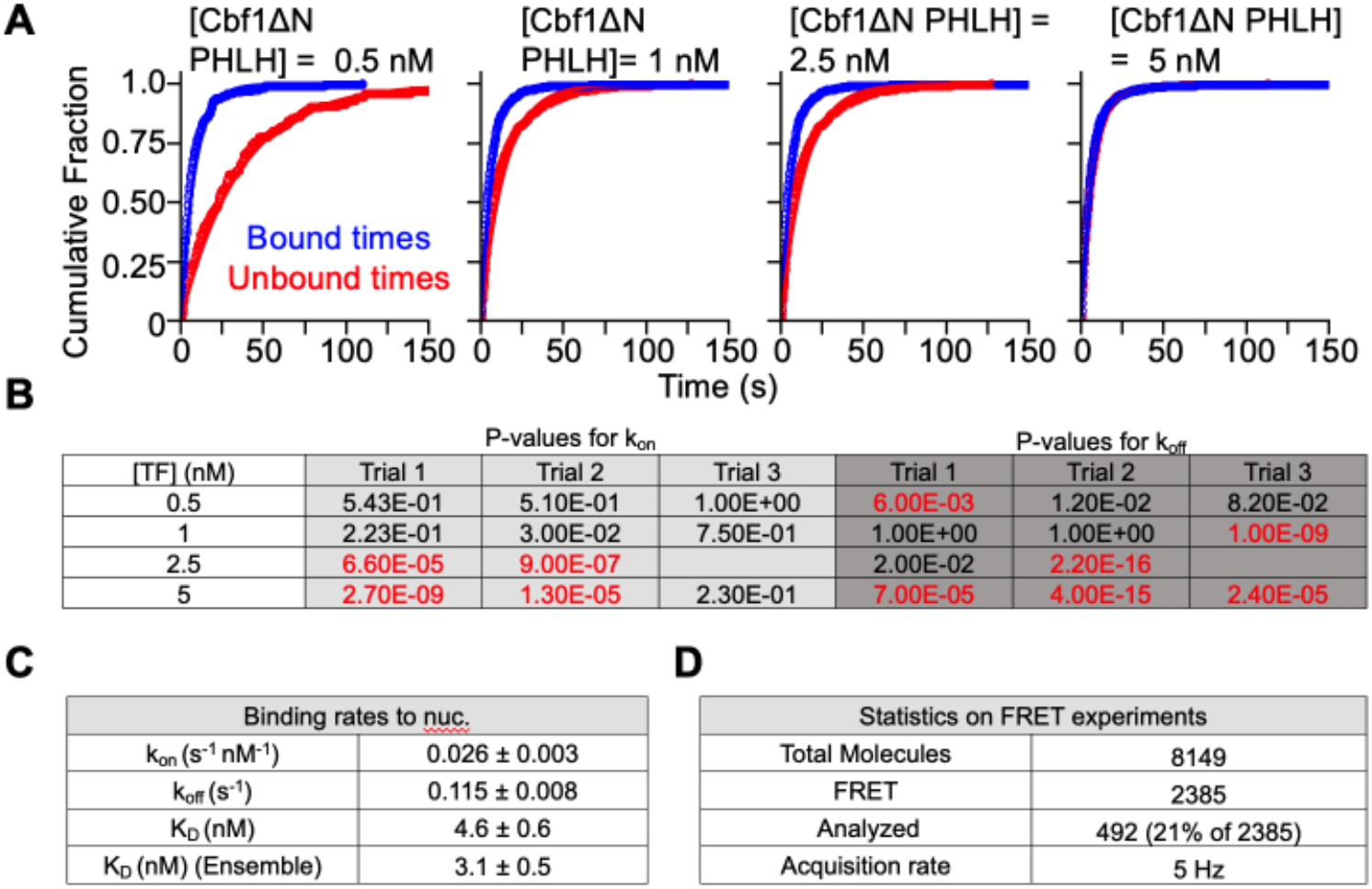
Single molecule characterization of Cbf1ΔN PHLH binding to Nucleosomes. (**A**) Cumulative sum distributions of bound (blue) and unbound dwell times (red) at all concentrations assayed. (**B**) Log-likelihood ratio tests were performed to determine whether to fit dwell time distributions to either a single or double exponential distribution. For each test, the null hypothesis was that the distribution follows a single-exponential decay. The null-hypothesis was rejected for P-values below 0.01. Rejected tests are indicated in red. Dwell time distributions were fit to the double-exponential model if 2 out of 3 replicates for all TF concentrations used in this experiment produced a P-value below 0.01. In this experiment, both the bound time and unbound time cumulative sums fit best to a single exponential. The resulting fits are included in (A) (**C**) The binding rate constant and dissociation rate determined from increasing concentrations of TF. (**D**) Statistics on the number of total molecules assayed. “FRET” refers to the number of molecules that emit from both Cy3 and a Cy5 fluorophores upon excitation at 532 nm and undergo fluctuations in FRET efficiency.

**Figure S15:**
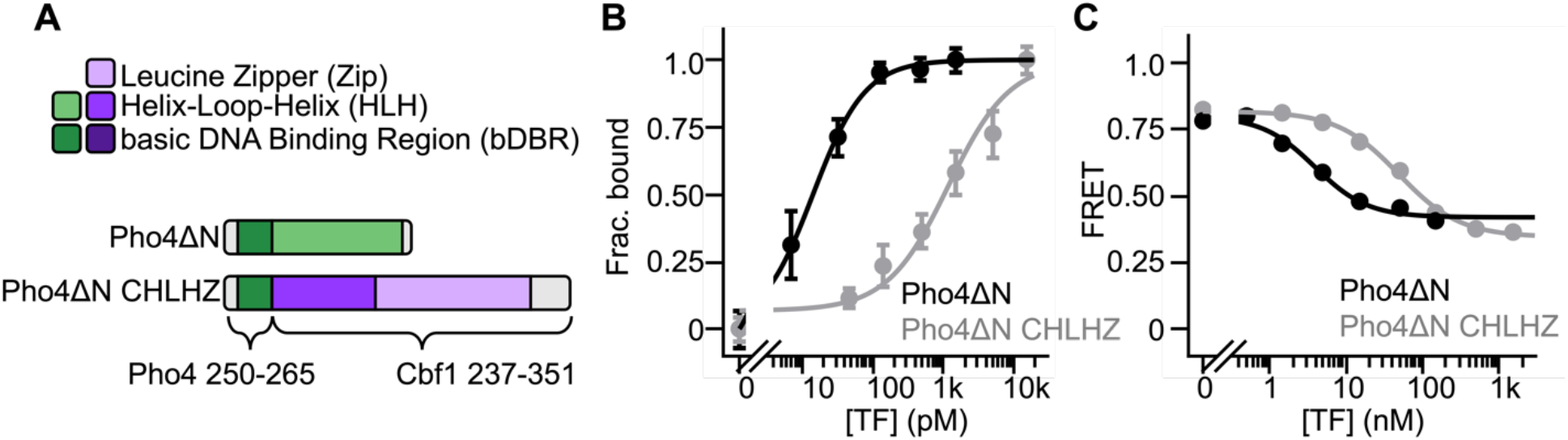
Ensemble fluorescence characterization of Pho4ΔN CHLHZ. (**A**) Domain diagrams of Pho4ΔN and Pho4ΔN CHLHZ in which the Cbf1 dimerization domain (Cbf1 residues 237-351) is inserted after Pho4 residues 250-265. (**B**) Ensemble PIFE measurements of Pho4ΔN and Pho4ΔN CHLHZ binding at increasing concentrations to DNA (S_1/2 Pho4ΔN DNA_ = 14 ± 1 pM, S_1/2 Pho4ΔN CHLHZ DNA_ = 1.2 ± 0.5 nM). The x-axis represents the estimated concentration of unbound protein. (**C**) Measuring binding to nucleosomes at increasing concentrations of Pho4ΔN and Pho4ΔN CHLHZ by the ensemble FRET assay (S_1/2 Pho4ΔN Nuc_ = 3.9 ± 0.9 nM, S_1/2 Pho4ΔN CHLHZ Nuc_ = 49 ± 6 nM).

**Figure S16.**
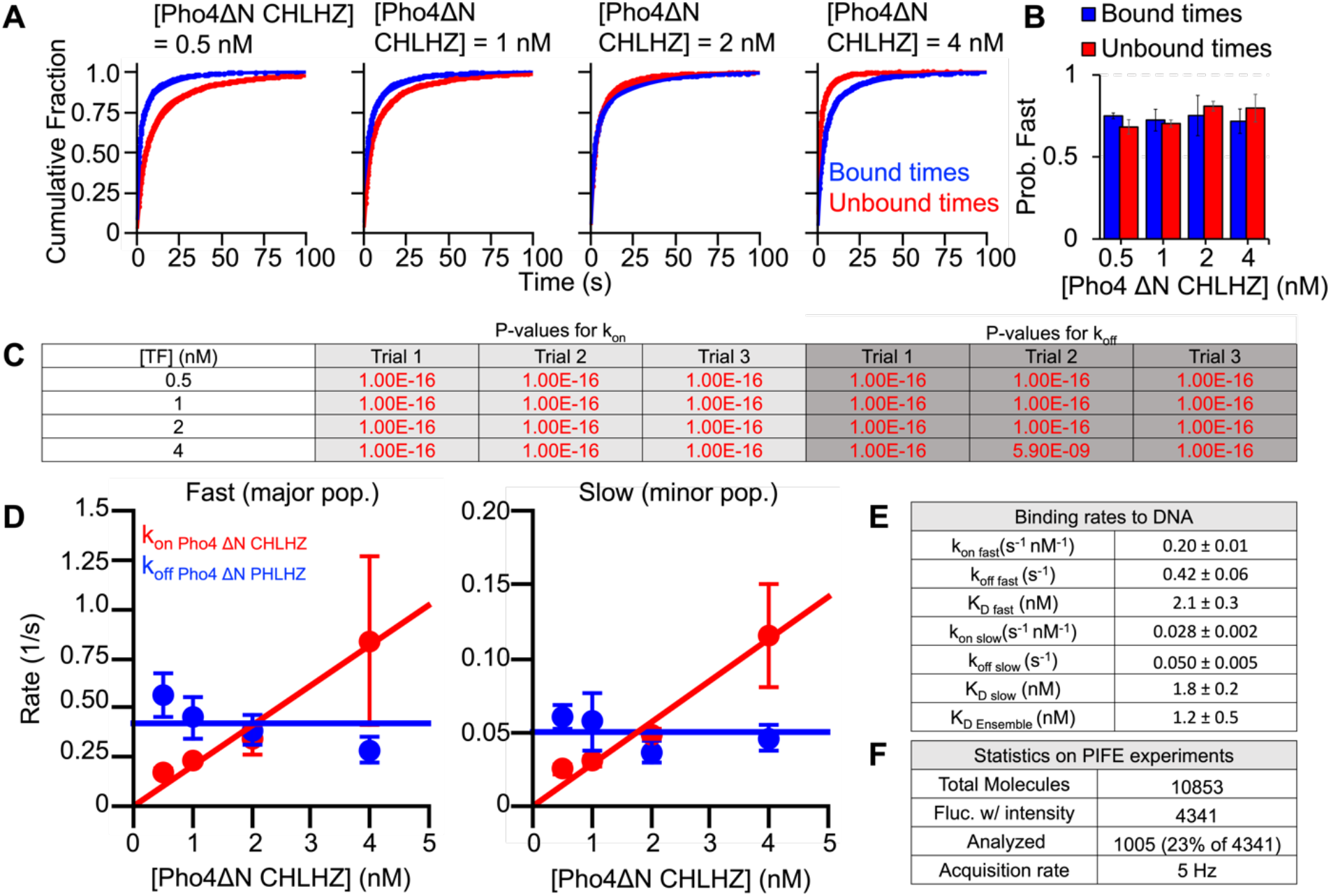
Single molecule characterization of Pho4ΔN CHLHZ binding to DNA. (**A**) Cumulative sum distributions of bound (blue) and unbound dwell times (red) at all concentrations assayed. (**B**) The probability of bound (blue bars) or unbound (red bars) dwell times belonging to the fast population for all concentrations assayed. (**C**) Log-likelihood ratio tests were performed to determine whether to fit dwell time distributions to either a single or double exponential distribution. For each test, the null hypothesis was that the distribution follows a single-exponential decay. The null-hypothesis was rejected for P-values below 0.01. Rejected tests are indicated in red. Dwell time distributions were fit to the double-exponential model if 2 out of 3 replicates for all TF concentrations used in this experiment produced a P-value below 0.01. In this experiment, both the bound time and unbound time cumulative sums fit best to a double exponential. The resulting fits are included in (A). (**D**) Major (left) and minor (right) binding and dissociation rates for increasing concentrations of TF. (**E**) Overall binding rate constants and dissociation rates determined from the fits in (D). (**F**) Statistics on the total number of molecules assayed. “Fluctuate with intensity” refers to the number of molecules with intensity values below 600 which was the empirically determined optimal intensity range for single-molecule PIFE.

**Figure S17.**
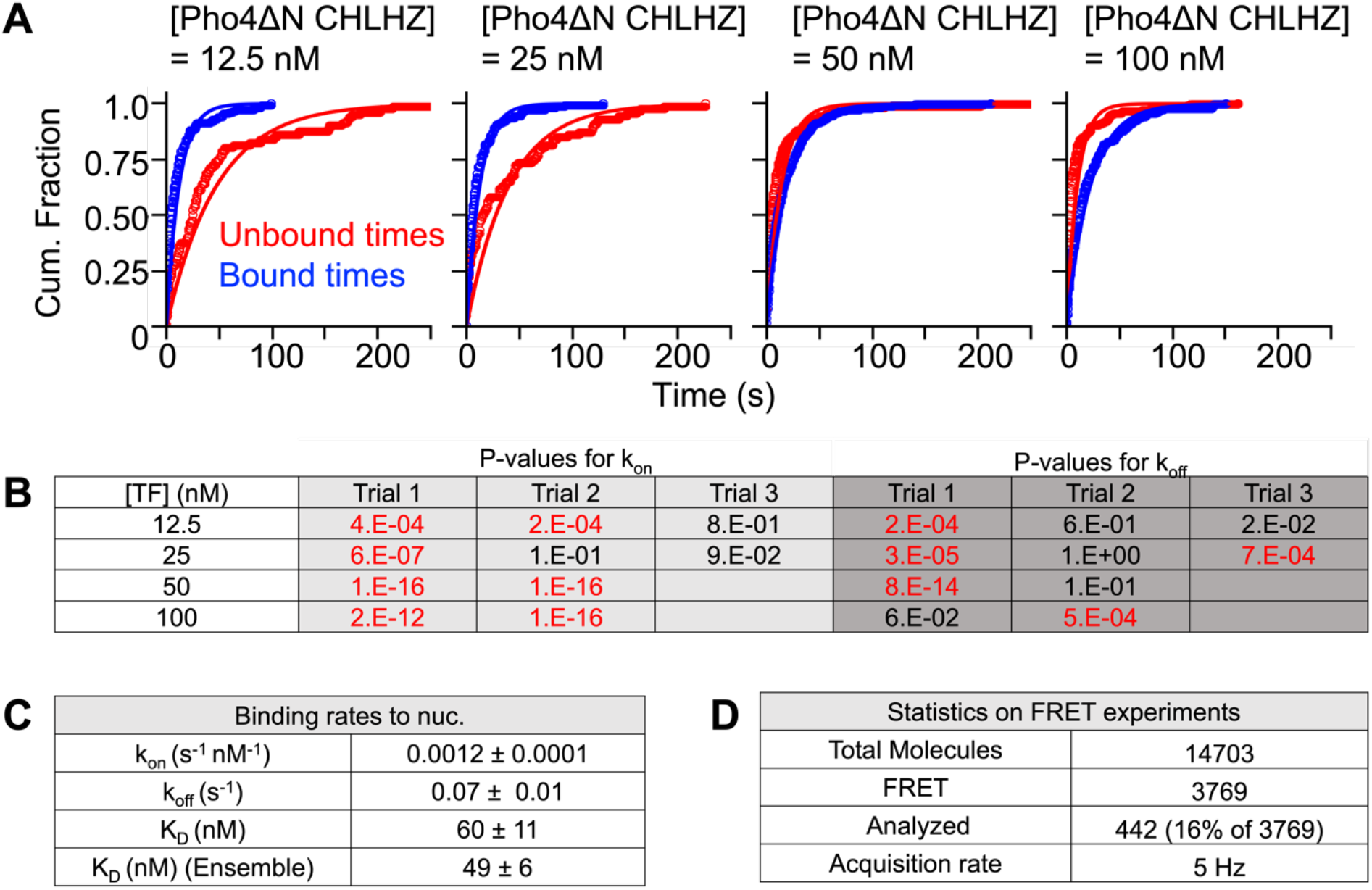
Single molecule characterization of Pho4ΔN CHLHZ binding to Nucleosomes. (**A**) Cumulative sum distributions of bound (blue) and unbound dwell times (red) at all concentrations assayed. (**B**) Log-likelihood ratio tests were performed to determine whether to fit dwell time distributions to either a single or double exponential distribution. For each test, the null hypothesis was that the distribution follows a single-exponential decay. The null-hypothesis was rejected for P-values below 0.01. Rejected tests are indicated in red. Dwell time distributions were fit to the double-exponential model if 2 out of 3 replicates for all TF concentrations used in this experiment produced a P-value below 0.01. In this experiment, both the bound time and unbound time cumulative sums fit best to a single exponential. The resulting fits are included in (A) (**C**) The binding rate constant and dissociation rate determined from increasing concentrations of TF. (**D**) Statistics on the number of total molecules assayed. “FRET” refers to the number of molecules that emit from both Cy3 and a Cy5 fluorophores upon excitation at 532 nm and undergo fluctuations in FRET efficiency.

**Figure S18.**
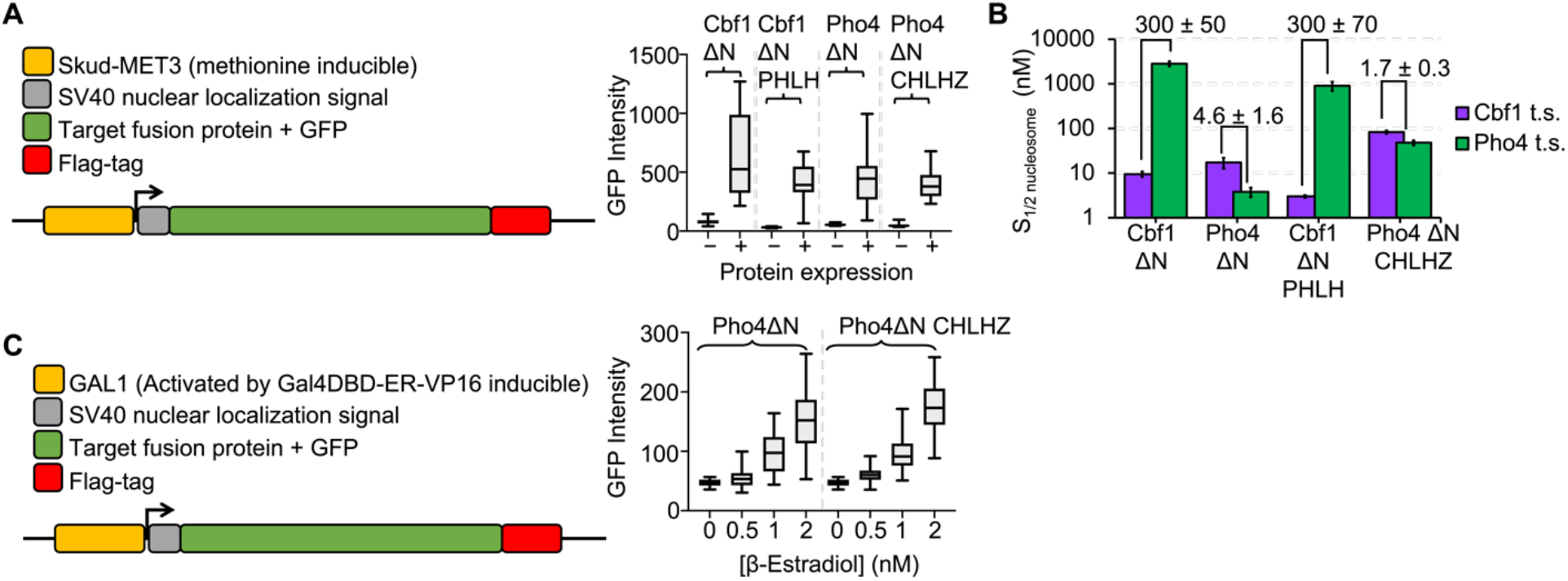
Constructs for in vivo expression of Pho4-Cbf1 chimeras. (**A**) GFP-tagged proteins are expressed from Met3 promoter in the absence of methionine. Protein expression is gauged by GFP intensity. Upon methionine depletion, all proteins are expressed to similar levels (right panel). (**B**) Ensemble FRET efficiency measurements of Cbf1ΔN, Cbf1ΔN PHLH, Pho4ΔN, and Pho4ΔN CHLHZ binding to the Cbf1 target site (t.s) (GGTCACGTGACC) and the Pho4 t.s. (CCCACGTGGG). S_1/2_ for each titration was determined from fitting to a binding isotherm: Cbf1ΔN: S_1/2 Cbf1 b.s._ = 9.3 ± 1.3 nM, S_1/2 Pho4 b.s._ = 2800 ± 300 nM; Cbf1 ΔN PHLH: S_1/2 Cbf1 b.s._= 3.0 ± 0.2 nM, S_1/2 Pho4 b.s._= 900 ± 210 nM; Pho4 ΔN: S_1/2 Cbf1 b.s._= 17.3 ± 4.7 nM, S_1/2 Pho4 b.s._= 3.8 ± 0.9; Pho4 ΔN CHLHZ: S_1/2 Cbf1 b.s._ = 83 ± 7 nM, S_1/2 Pho4 b.s._= 48 ± 6 nM. Bar graph represents S_1/2_ values from the ensemble FRET measurements. Proteins containing the Pho4 basic region (Pho4ΔN and Pho4ΔN CHLHZ) bind similarly to both sites. Proteins containing the Cbf1 basic region (Cbf1ΔN and Cbf1ΔN PHLH) have a strong preference for the Cbf1 consensus site. (**C)** High regulation of protein expression levels was accomplished using a GAL1 promoter activated by GAL4DBD-ER-VP16. Titration of β-Estradiol shows concentration-dependent expression of GFP intensity and that both proteins are expressed to similar levels (right panel). For both plots, whiskers represent the complete range of GFP intensity values (minimum to maximum).

**Figure S19.**
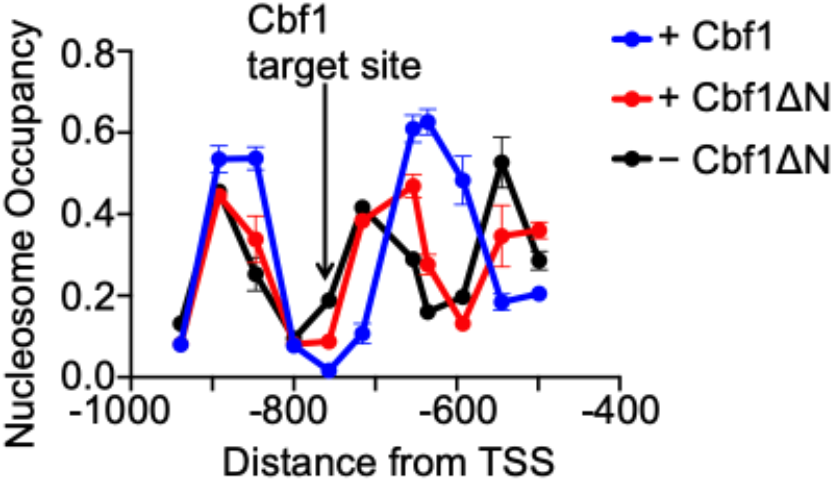
Nucleosome occupancy in the HO promoter from nucleosome −5 to nucleosome -3. The Cbf1 target site is placed 43 bp from the nucleosome dyad in the -4 nucleosome. Expression of either Cbf1 or Cbf1ΔN results in a shift of the −4 nucleosome away from the Cbf1 target site. Full-length Cbf1 produces a much larger NDR. Cbf1 data is from (Yan et al., 2018).

## SUPPLEMENTAL TABLES

**Table S1.**
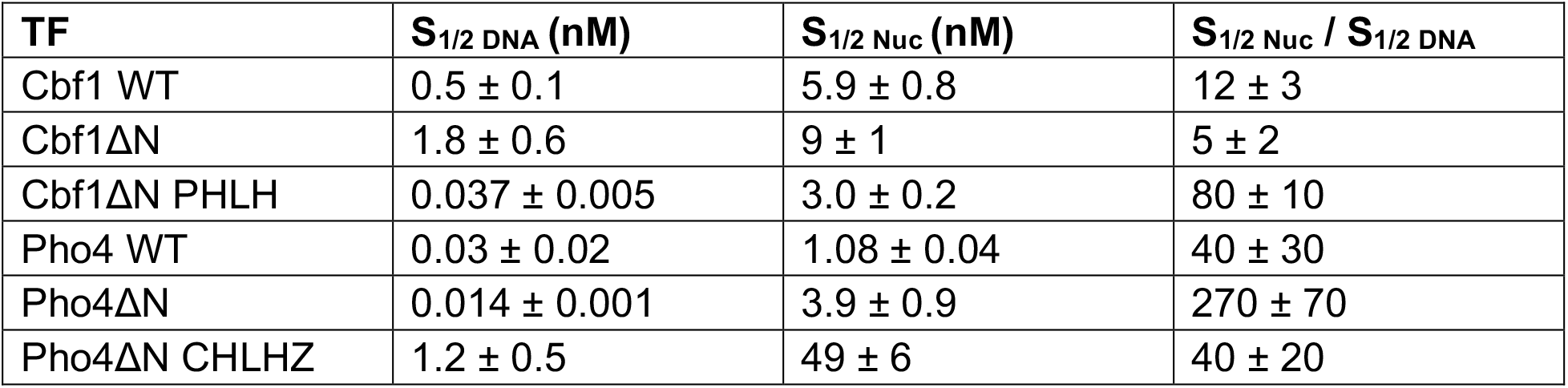
Summary of binding affinities determined by ensemble fluorescence measurements

**Table S2.** Summary of primary binding and dissociation kinetics to and from DNA determined by smPIFE measurements

| TF | $k_{\text{on DNA}} \text{ (s}^{-1} \text{ nM}^{-1}\text{)}$ | $k_{\text{off DN}} \text{ (s}^{-1}\text{)}$ | $K_D \text{ (nM)}$ |
| --- | --- | --- | --- |
| Cbf1 WT | $0.025 \pm 0.006$ | $0.30 \pm 0.05$ | $12.0 \pm 3.5$ |
| Cbf1ΔN | $0.16 \pm 0.01$ | $0.39 \pm 0.02$ | $2.4 \pm 0.2$ |
| Cbf1ΔN PHLH | $9.8 \pm 1.3$ | $0.32 \pm 0.04$ | $0.032 \pm 0.006$ |
| Pho4 WT | $22 \pm 5$ | $0.31 \pm 0.04$ | $0.014 \pm 0.003$ |
| Pho4ΔN | $40 \pm 10$ | $0.33 \pm 0.01$ | $0.008 \pm 0.002$ |
| Pho4ΔN CHLHZ | $0.20 \pm 0.01$ | $0.42 \pm 0.06$ | $2.1 \pm 0.3$ |

**Table S3.** Summary of primary binding and dissociation kinetics to and from nucleosomes determined by smFRET measurements

| TF | $k_{\text{on Nuc}} \text{ (s}^{-1} \text{ nM}^{-1}\text{)}$ | $k_{\text{off Nuc}} \text{ (s}^{-1}\text{)}$ | $K_D \text{ Nuc (nM)}$ |
| --- | --- | --- | --- |
| Cbf1 WT | $2.10\text{E-}04 \pm 2.00\text{E-}05$ | $1.11\text{E-}02 \pm 7.00\text{E-}04$ | $53 \pm 6$ |
| Cbf1ΔN | $6.12\text{E-}04 \pm 1.00\text{E-}04$ | $5.07\text{E-}03 \pm 5.78\text{E-}04$ | $8.3 \pm 1.7$ |
| Cbf1ΔN PHLH | $2.69\text{E-}02 \pm 3.07\text{E-}03$ | $1.15\text{E-}01 \pm 8.27\text{E-}03$ | $4.3 \pm 0.6$ |
| Pho4 WT | $2.00\text{E-}01 \pm 4.00\text{E-}02$ | $3.00\text{E-}01 \pm 2.00\text{E-}02$ | $1.5 \pm 0.3$ |
| Pho4ΔN | $1.80\text{E-}01 \pm 2.23\text{E-}02$ | $5.93\text{E-}01 \pm 3.23\text{E-}02$ | $3.3 \pm 0.4$ |
| Pho4ΔN CHLHZ | $1.15\text{E-}03 \pm 8.30\text{E-}05$ | $6.88\text{E-}02 \pm 1.16\text{E-}02$ | $60 \pm 11$ |

**Table S4.** Summary of nucleosome binding and dissociation kinetics relative to DNA.

| TF | $K_{on} \text{ Nuc} / K_{on} \text{ DNA}$ | $K_{off} \text{ Nuc} / K_{off} \text{ DNA}$ | $K_D \text{ Nuc} / K_D \text{ DNA}$ |
| --- | --- | --- | --- |
| Cbf1 WT | $0.008 \pm 0.002$ | $0.037 \pm 0.007$ | $4.4 \pm 1.4$ |
| Cbf1ΔN | $0.0038 \pm 0.0007$ | $0.013 \pm 0.002$ | $3.4 \pm 0.7$ |
| Cbf1ΔN PHLH | $0.0027 \pm 0.0005$ | $0.36 \pm 0.05$ | $130 \pm 30$ |
| Pho4 WT | $0.009 \pm 0.003$ | $1.0 \pm 0.1$ | $110 \pm 40$ |
| Pho4ΔN | $0.004 \pm 0.001$ | $1.8 \pm 0.1$ | $400 \pm 100$ |
| Pho4ΔN CHLHZ | $0.0058 \pm 0.0005$ | $0.16 \pm 0.04$ | $29 \pm 7$ |

**Table S5.** Yeast strains used in this study.

| Strain name | Genotype | Notes |
| --- | --- | --- |
| yHC54-Cbf1dN | *can1, his3, leu2, trp1, ura2, hoprurs2 (nt-1298 to -272), ADE2, cbf1:GAL1pr-CBF1:TRP1, CLN2:HOPr-swi5mut-43Pho4-GFP:URA3, his3:SkudMET3pr-CBF1dN GFP 3xFLAG:HIS3* | Cbf1 binding motif and Cbf1dN driven by Met promoter |
| yHC54-Cbf1dN_PDD | *can1, his3, leu2, trp1, ura2, hoprurs2 (nt-1298 to -272), ADE2, cbf1:GAL1pr-CBF1:TRP1, CLN2:HOPr-swi5mut-43Pho4-GFP:URA3, his3:SkudMET3pr-CBF1dN_PDD GFP 3xFLAG:HIS3* | Cbf1 binding motif and Cbf1dN_PDD driven by Met promoter |
| yHC54-Pho4dN | *can1, his3, leu2, trp1, ura2, hoprurs2 (nt-1298 to -272), ADE2, cbf1:GAL1pr-CBF1:TRP1, CLN2:HOPr-swi5mut-43Pho4-GFP:URA3, his3:SkudMET3pr-PHO4dN GFP 3xFLAG:HIS3* | Cbf1 binding motif and Pho4dN driven by Met promoter |
| yHC54-Pho4dN_CDD | *can1, his3, leu2, trp1, ura2, hoprurs2 (nt-1298 to -272), ADE2, cbf1:GAL1pr-CBF1:TRP1, CLN2:HOPr-swi5mut-43Pho4-GFP:URA3, his3:SkudMET3pr-PHO4dN_CDD GFP 3xFLAG:HIS3* | Cbf1 binding motif and Pho4dN_CDD driven by Met promoter |
| yHC55-Cbf1dN | *can1, his3, leu2, trp1, ura2, hoprurs2 (nt-1298 to -272), ADE2, cbf1:GAL1pr-CBF1:TRP1, CLN2:HOPr-swi5mut-43Cbf1-GFP:URA3, his3:SkudMET3pr-CBF1dN GFP 3xFLAG:HIS3* | Pho4 binding motif and Cbf1dN driven by Met promoter |
| yHC55-Cbf1dN_PDD | *can1, his3, leu2, trp1, ura2, hoprurs2 (nt-1298 to -272), ADE2, cbf1:GAL1pr-CBF1:TRP1, CLN2:HOPr-swi5mut-43Cbf1-GFP:URA3, his3:SkudMET3pr-CBF1dN_PDD GFP 3xFLAG:HIS3* | Pho4 binding motif and Cbf1dN_PDD driven by Met promoter |
| yHC55-Pho4dN | *can1, his3, leu2, trp1, ura2, hoprurs2 (nt-1298 to -272), ADE2, cbf1:GAL1pr-CBF1:TRP1, CLN2:HOPr-swi5mut-43Cbf1-* | Pho4 binding motif and Pho4dN driven by Met promoter |
|  | *GFP:URA3, his3:SkudMET3pr-*<br>*PHO4dN_GFP_3xFLAG:HIS3* |  |
| yHC55-Pho4dN_CDD | *can1, his3, leu2, trp1, ura2, hoprurs2 (nt-*<br>*1298 to -272), ADE2, cbf1:GAL1pr-*<br>*CBF1:TRP1, CLN2:HOPr-swi5mut-43Cbf1-*<br>*GFP:URA3, his3:SkudMET3pr-*<br>*PHO4dN_CDD_GFP_3xFLAG:HIS3* | Pho4 binding motif and<br>Pho4dN_CDD driven by<br>Met promoter |
| yHC64-Pho4dN | *can1, his3, leu2, trp1, ura2, hoprurs2 (nt-*<br>*1298 to -272), ADE2, cbf1:GAL1pr-*<br>*CBF1:TRP1, CLN2:HOPr-swi5mut-43Pho4-*<br>*GFP:URA3, his3:ADH1pr-GAL4DBD-ER-*<br>*VP16::GAL1pr_-*<br>*PHO4dN_GFP_3xFLAG:HIS3* | Pho4 binding motif and<br>Pho4dN driven by GAL1<br>promoter |
| yHC64-Pho4dN_CDD | *can1, his3, leu2, trp1, ura2, hoprurs2 (nt-*<br>*1298 to -272), ADE2, cbf1:GAL1pr-*<br>*CBF1:TRP1, CLN2:HOPr-swi5mut-43Pho4-*<br>*GFP:URA3, his3:ADH1pr-GAL4DBD-ER-*<br>*VP16::GAL1pr-*<br>*PHO4dN_CDD_GFP_3xFLAG:HIS3* | Pho4 binding motif and<br>Pho4dN_CDD driven by<br>GAL1 promoter |

**Table S6.**
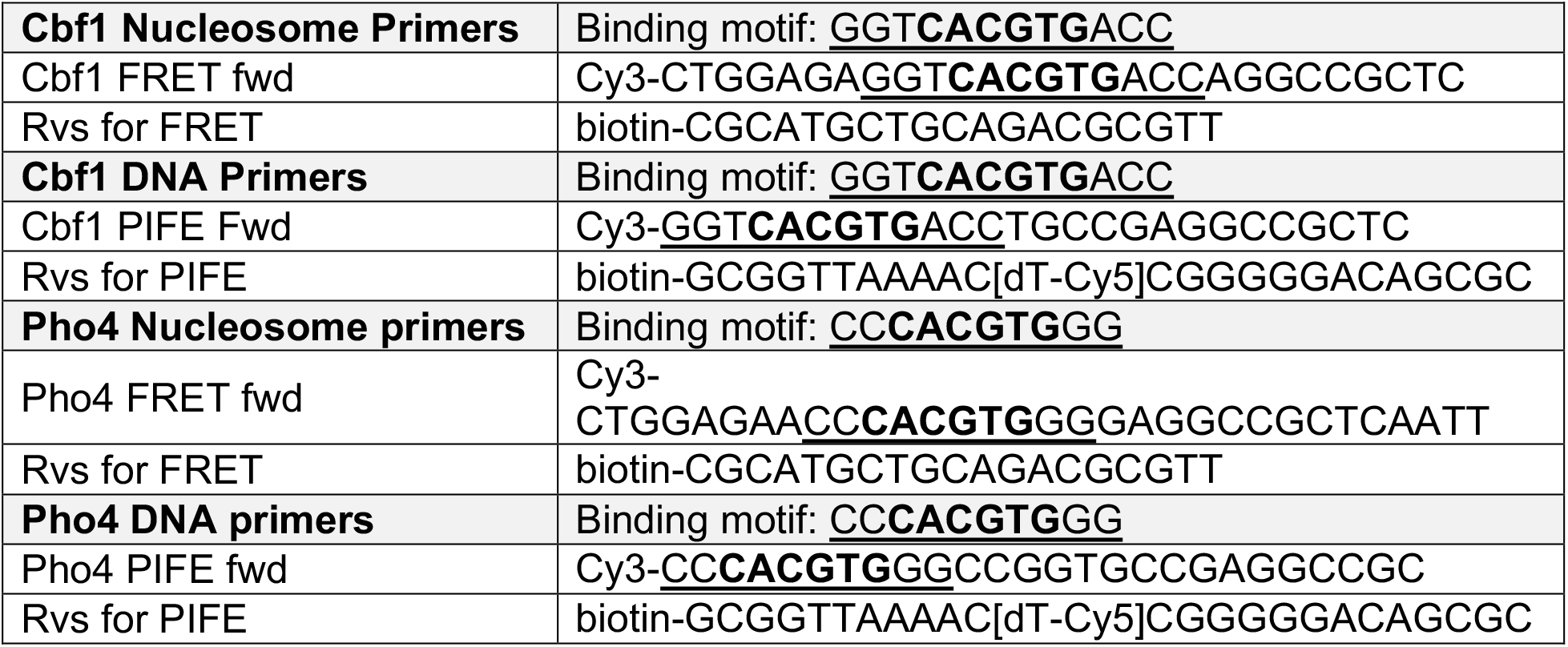
PCR primers for DNA used in single-molecule studies

**Table S7.**
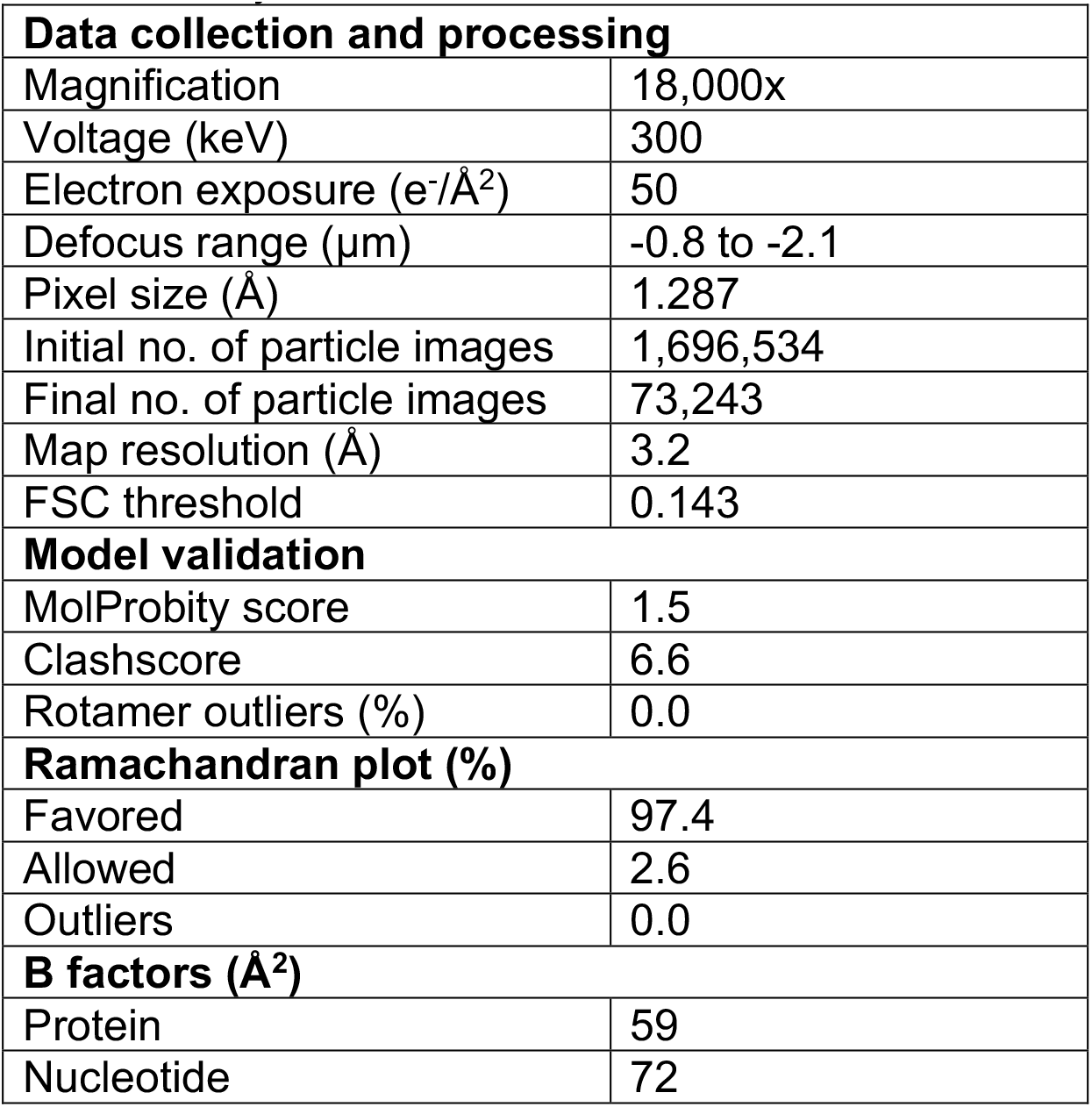
Cryo-EM data collection and refinement statistics

| **Data collection and processing** |  |
| --- | --- |
| Magnification | 18,000x |
| Voltage (keV) | 300 |
| Electron exposure (e<sup>-</sup>/Å<sup>2</sup>) | 50 |
| Defocus range (μm) | -0.8 to -2.1 |
| Pixel size (Å) | 1.287 |
| Initial no. of particle images | 1,696,534 |
| Final no. of particle images | 73,243 |
| Map resolution (Å) | 3.2 |
| FSC threshold | 0.143 |
| **Model validation** |  |
| MolProbity score | 1.5 |
| Clashscore | 6.6 |
| Rotamer outliers (%) | 0.0 |
| **Ramachandran plot (%)** |  |
| Favored | 97.4 |
| Allowed | 2.6 |
| Outliers | 0.0 |
| **B factors (Å<sup>2</sup>)** |  |
| Protein | 59 |
| Nucleotide | 72 |

**Table S8.**
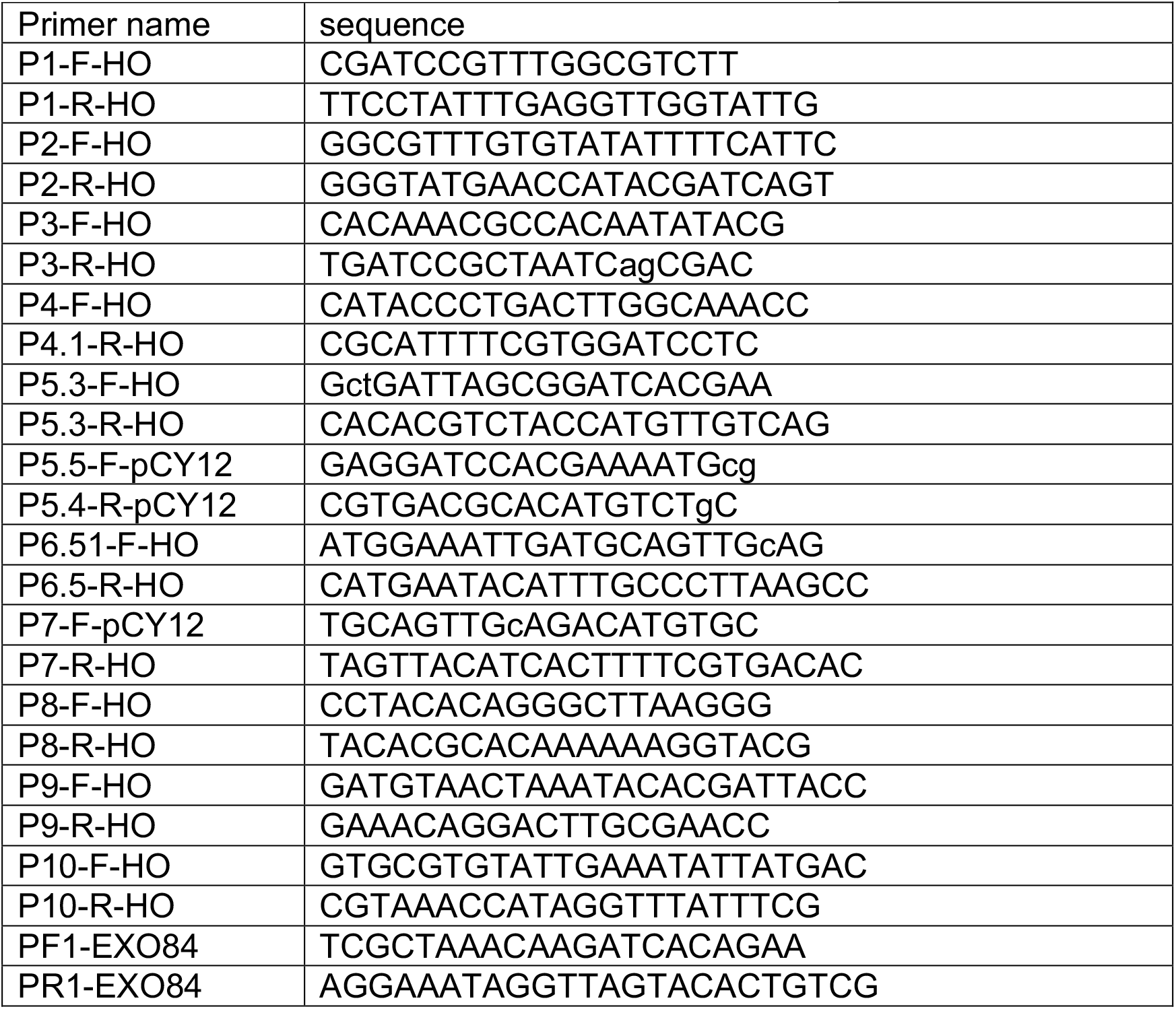
MNase-qPCR primers used in this study

